# Emergence of an adaptive epigenetic cell state in human bladder urothelial carcinoma evolution

**DOI:** 10.1101/2021.10.30.466556

**Authors:** Yu Xiao, Wan Jin, Kaiyu Qian, Kai Wu, Gang Wang, Wei Jiang, Rui Cao, Lingao Ju, Yan Zhao, Hang Zheng, Tongzu Liu, Luyuan Chang, Zilin Xu, Ting Wang, Jun Luo, Liuying Shan, Fang Yu, Xintong Chen, Dongmei Liu, Hong Cao, Zhonghua Yang, Sheng Li, Hongjie Shi, Zhongqiang Guo, Yan Gong, Nan Liu, Shenjuan Li, Yejinpeng Wang, Xinyue Cao, Wenjun Ding, Wei Zhou, Diansheng Cui, Ye Tian, Chundong Ji, Yongwen Luo, Xin Hong, Haoli Ma, Fangjin Chen, Minsheng Peng, Yi Zhang, Xinghuan Wang

**Author notes:** Corresponding authors: Dr. Xinghuan Wang. These authors contributed equally to this work (Yu Xiao, Wan Jin, Kaiyu Qian, Kai Wu, Gang Wang, Wei Jiang, Rui Cao, Lingao Ju, Yan Zhao, Hang Zheng).

## Abstract

Intratumor heterogeneity (ITH) of bladder cancer (BLCA) facilitates therapy resistance and immune evasion to affect clinical prognosis directly. However, the molecular and cellular mechanism generating ITH in BLCA remains elusive. Here we show that a TM4SF1-positive cancer subpopulation (TPCS) drives ITH diversification in BLCA. By extensive profiling of the epigenome and transcriptome of BLCA from 79 donors across all stages, we elucidated the evolution trajectories of luminal and basal BLCA. TPCS emerges from the basal trajectory and shows extensive transcriptional plasticity with a distinct epigenomic landscape. Clinically, TPCS were enriched in advanced stage patients and associated with poor prognosis. Our results showed how cancer adapts to its environment by adopting a stem cell-like epigenomic landscape.

## Introduction

Bladder urothelial carcinoma (BLCA), which causes 212,500 deaths worldwide annually^1^, is a biologically diverse ensemble. The phenotypic diversity, in other words, intratumor heterogeneity (ITH) of BLCA determined their biological behaviour. ITH correlates with clinical stages, mutation burdens, and DNA copy number changes^2^ and less favourable survival outcome in both BLCA^3^ and other cancer types^4–7^. Whilst previous studies in other cancer types suggested that epigenetic plasticity^8–13^ and genomic mutation^9, 10, 14^ lead to ITH, the exact molecular mechanism generating ITH in BLCA remains elusive. Clinically, BLCA are classified into two distinct categories: the less malignant non-muscle-invasive BLCA (NMIBC), including Ta, T1 and/or CIS (Tis) stages, and muscle-invasive BLCA (MIBC) with rapid growth rate and metastasis potential^14–19^. Studies on rodent models have classified BLCA into luminal-type and basal-type cancers by cell-of-origins^20, 21^, which resemble human NMIBC and MIBC, respectively^22^. The fact that basal- and luminal-like phenotypes might co-exist in the same tumour or different urinary tract tumours of a single individual^9, 23^ suggests interconvertibility between the classes. Therefore, the cellular-origin, genetic diversity, and epigenetic heterogeneity might all contribute to ITH in BLCA.

## Results

### Single cell landscape of normal bladder and BLCA

To characterize BLCA evolution from the single cell (sc) level (Supplementary Fig. 1), we performed scRNA and scATAC sequencing on 56 tissues from 26 donors, including 23 BLCA patients and 3 Donations after Cardiac Death (DCD) (Supplementary Table 1-4). Also, scRNA data of 11 BLCA tissues from the public domain^24^ was included. After stringent quality control^25–27^, 133,953 cells remained, including epithelia, endothelia, fibroblasts, myeloid cells, CD3^+^ T-lymphocytes, B-lymphocytes, and NK cells (Fig. 1, Supplementary Table 3-6). These cells were derived from different donors (Fig. 1C) and different tissue types (Fig. 1D).

**Figure 1.**
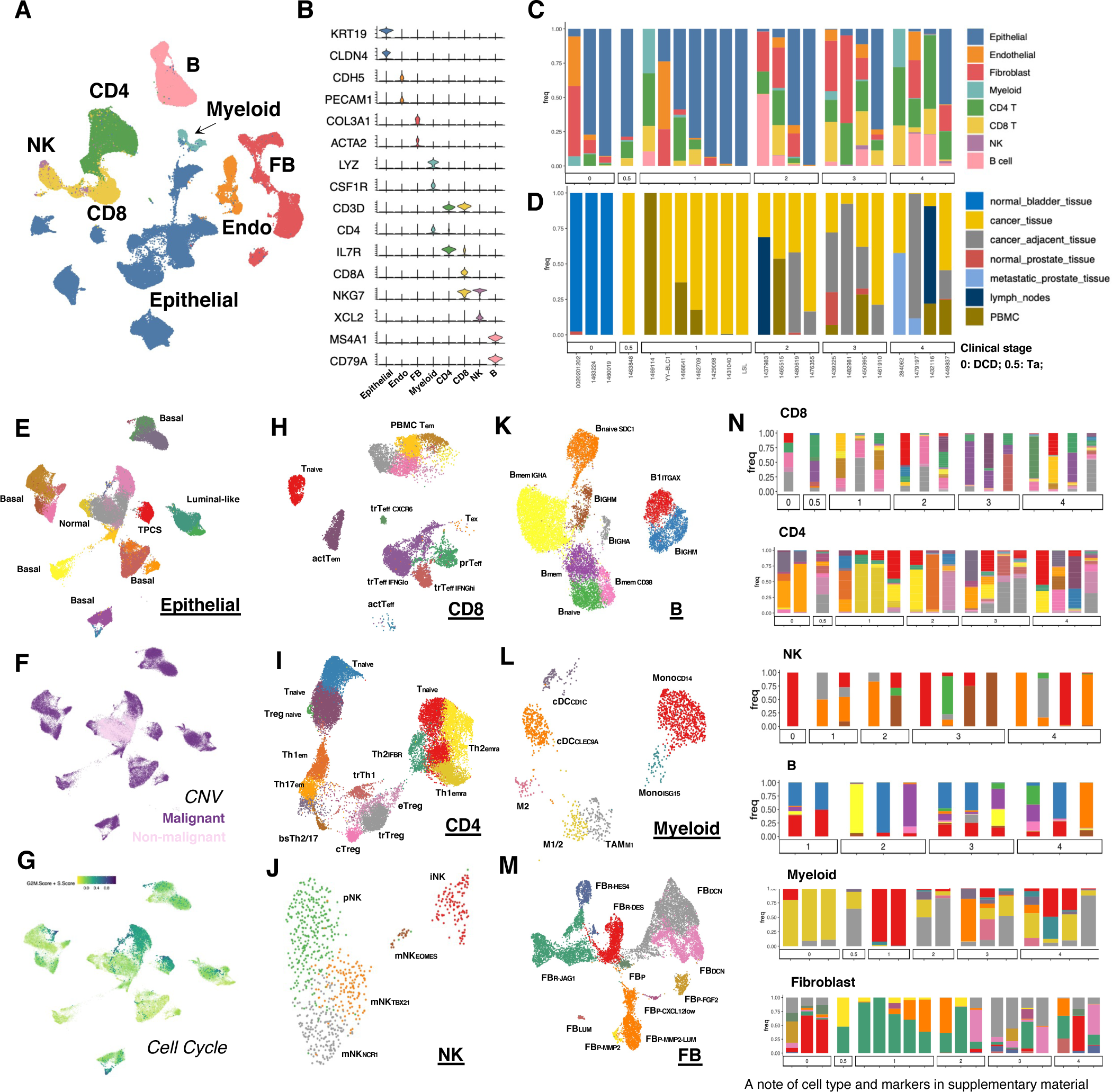
Single cell landscape of BLCA. **(A)** UMAP-projection of scRNA sequencing from bladder tissues. Epithelial cells, fibroblast (FB), endothelial cells (Endo), macrophage and monocyte (Myeloid), CD4^+^ helper T cells (CD4), CD8^+^ cytotoxic T cells (CD8), natural killer cells (NK), and B lymphoid cells (B) were shown. **(B)** Marker gene expression for each cell type. **(C)** Cell type composition in bladder tissues of in-house sequenced sample donors. Donors were separated by T stage (0: healthy organ donor; 0.5: Ta; 1-4: T1-T4; Ta/T1: NMIBC; T2-T4: MIBC). **(D)** Tissue type prevalence for each donor. Donors were separated by clinical stage as in (C). **(E)** UMAP-projection of epithelial cells. Normal epithelial cells were identified by inferred CNV in (F). Fast-cycling, proliferating cancer cells were identified by cell cycle score (G). Luminal-like, basal-like, and TM4SF1-positive cancer cells were identified by marker gene expression (Supplementary Fig. 2). **(F)** Inferred CNV from integrated scATAC data. **(G)** Inferred cell cycle score. **(H-M)** UMAP-projection of CD8-positive cytotoxic T, CD4-positive helper T, NK, B, Myeloid and Fibroblast (Supplementary Fig. 6-7 and 9-12). **(N)** Cell type prevalence in each donor. Bars are coloured as in panels (H)-(M), respectively. Donors are separated by clinical stage as (C). Significant changes in cellular composition through clinical stage advancements were found for CD8 (recruitment of bystander T_eff_), CD4 (emergence of tumour-reactive trT_reg_), and fibroblasts.

Epithelial cells dominate the population in cancer (Fig. 1E). Driven by donor-specific copy number variation (CNV), urothelium epithelia originated cancer cells segregated into donor-specific clusters, whilst CNV-free cells from tumour and normal urothelial tissues tended to cluster together (Fig. 1E-F). Luminal-like and basal-like cancer cells could be identified by the expression of *GATA3/FGFR3* or *KRT5/KRT6A/KRT7*, respectively (Fig. 1E and Supplementary Fig. 2). Two populations of cancer cells are marked with robust proliferating gene expression: a TM4SF1-positive cancer subpopulation (TPCS), and a TM4SF1-negative, fast proliferating cell population (FPC) (Fig. 1E-G and 3H-I). Both TPCS and FPC highly expressed cell-cycle related genes such as *H2AZ1* and *H2AZ2,* whilst TPCS additionally overexpressed TM4SF1, PI3, and a set of keratins (KRT7/KRT6A/KRT6B) (Fig. 3I and Supplementary Fig. 2).

We further characterized 1,163 normal epithelial cells from healthy organ donors in detail. These cells were classified into basal (Bs1/2/3), intermediate (Im1/2/3), umbrella (Um) by the expression of *SHH*, *KRT5* and uroplakins^28^ and a small number of urothelium stem/progenitor cell (BsP) (Fig. 2A-B). BsP is highly proliferative (Fig. 2C) and overexpressed *TM4SF1* (Fig. 3I). Basal cell lineage is clearly separated from the superficial cells not only by conventional markers *KRT5/UPK2/UPK3A/UPK3B* but also by *HES1* and *VEGFA* expression (Fig. 2D). Bs2/3 lineage are distinct from BsP/Bs1 by that they overexpressed *PCSK5* and *SGK1* and have no *HBEGF* expression (Fig. 2D), suggesting a functional heterogeneity between the basal cells.

**Figure 2.**
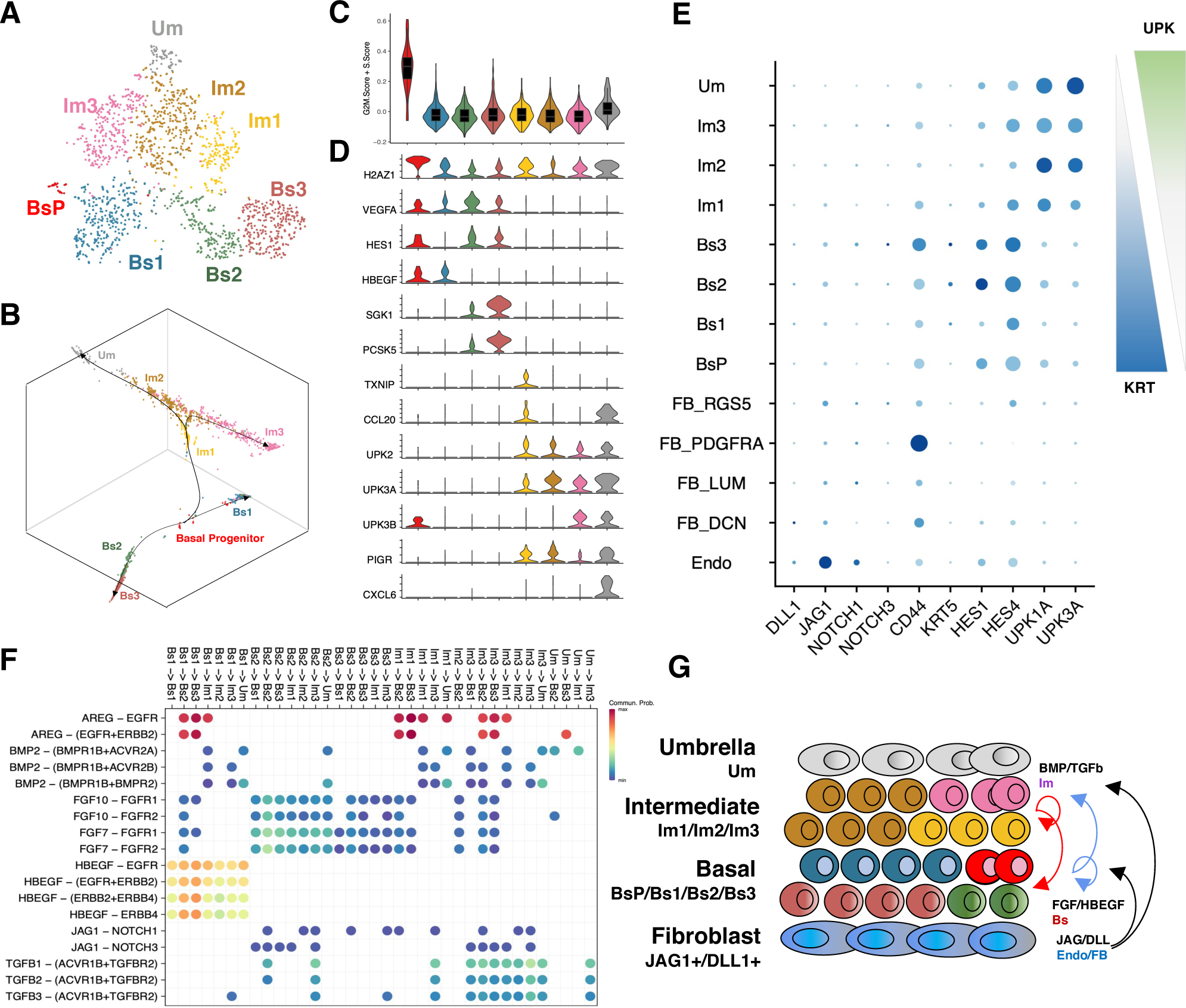
Bladder urothelium cell fate determination by mutual signalling. **(A)** TSNE-projection of normal bladder urothelium epithelial cells from the 3 healthy organ donors. BsP: basal progenitor cell (SHH^+^/CD44^+^/TM4SF1^+^); Bs1: basal cell 1 (IGFBP2^+^/KRT17^+^/PTGS2^+^); Bs2: basal cell 2 (HES1^+^/ELF3^+^); Bs3: basal cell (HES1^+^/PCSK5^+^); Im1: intermediate cell 1 (TXNIP^+^); Im2: intermediate cell 2 (MMP7^+^); Im3: intermediate cell 3 (GDF15^+^/KRT13^+^); Um: umbrella cell (CD24^+^/KRT18^+^). **(B)** Developmental trajectory of urothelium on 3D diffusion map projection. Basal progenitor cell evolves into three lineages: Bs1, Bs2/Bs3, and Im1/2/3->Um. **(C)** Cell cycle score of each cluster in (A). **(D)** Marker gene expression of each cluster in (A). Note that HES1/VEGFA are only positive in BsP/Bs1/Bs2, whilst UPK2/3A/3B and PIGR are only positive in intermediate/umbrella cells. Umbrella cell is identified by CXCL6 expression. **(E)** NA expression of basal/intermediate markers (KRT5/CD44 as basal, and UPK1A/3A as intermediate markers) and Notch signalling pathway ligand/receptors. Note that whilst Notch downstream effector HES1/HES4 are upregulated in basal cells, the main source of Notch ligand DLL1/JAG1 comes from the underlying endothelial/fibroblast population. **(F)** Cell-cell signalling probability between epithelial cells suggesting Bs1 provides HBEGF to all other cells, Bs2/Bs3 providing FGF, and intermediate cells contribute BMP/TGFβ ligands. **(G)** Anatomy of urothelium from the basal (bottom) side to the luminal (top) side shows cell types and the identified cell-cell signalling events between them. Basal/intermediate cell lineages are likely to be separated by Notch signalling activity strength, as basal cells are more proximal to the major Notch ligand provider.

**Figure 3.**
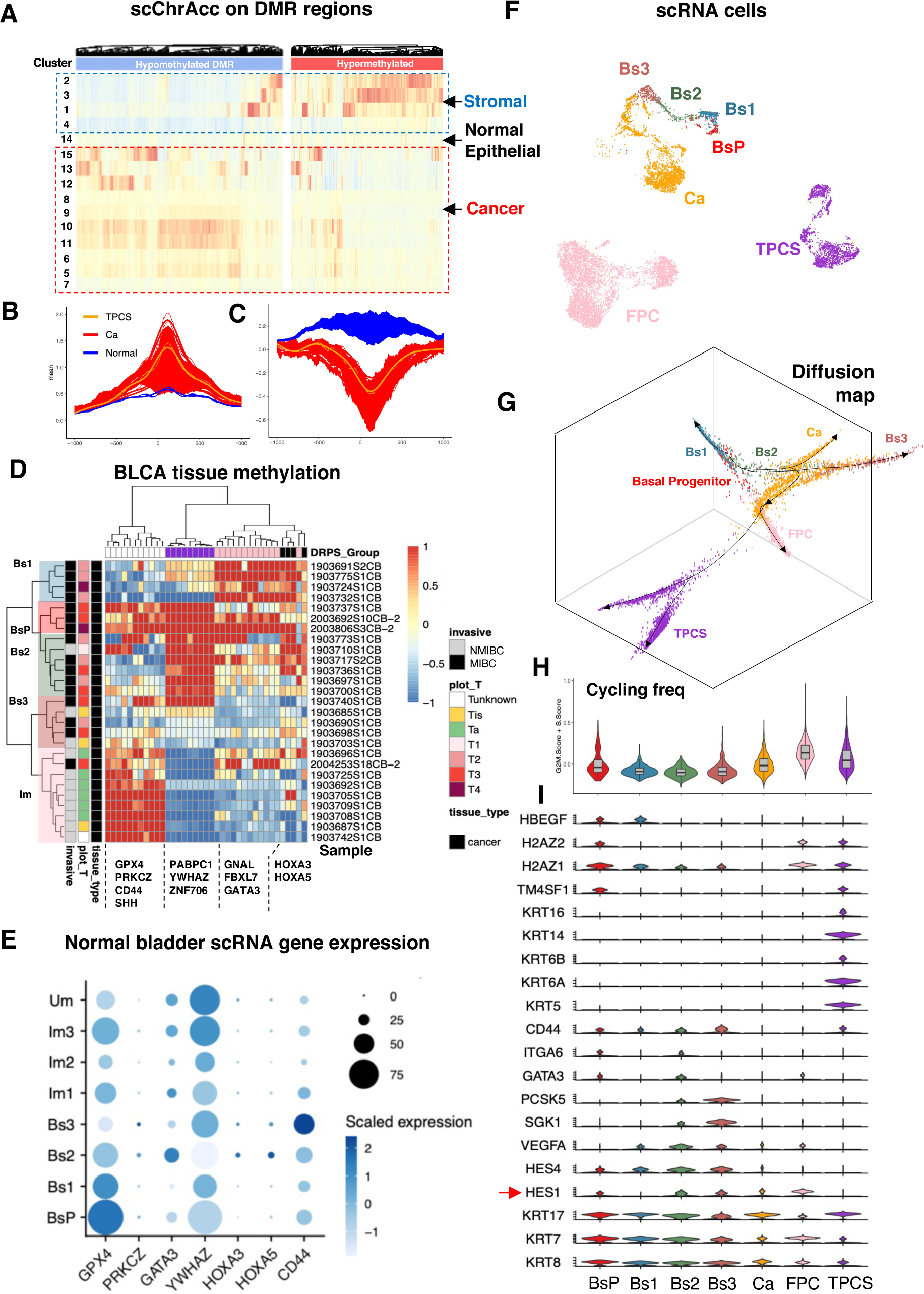
Origin of muscle-invasive and TM4SF1-positive bladder cancer cells in urothelium. **(A)** Single cell chromatin accessibility on MIBC-specific differentially methylated regions (DMR) among stromal (2/3/1/4), normal epithelial (14), and cancer (5-13 and 15). MIBC-hypo DMR are closed in normal epithelial cells but open in cancer. MIBC-hyper DMR are only open in stromal and normal epithelia. **(B-C)** Chromatin accessibility of MIBC-hyper DMR (B) and MIBC-hypo DMR (C) in normal epithelial, cancer and TPCS. **(D)** Unsupervised hierarchical clustering of DMR methylation level (Z-scaled) of BLCA tissue samples on selected hypermethylated DMR. Samples were labelled by invasiveness (Invasive) and clinical stage (T). NMIBC is clearly separated from MIBC by hypermethylation in GPX4/PRKCZ/SHH/CD44 genes, and MIBC could be classified into 4 clusters by DMR methylation level. **(E)** RNA expression of DMR-associated genes in different types of urothelial cells. **(F)** UMAP-projection of normal urothelium basal cells (BsP, Bs1, Bs2, Bs3), early-stage (CNV-low) BLCA cells (Ca), TM4SF1-positive cancer cells (TPCS), and fast-proliferating cancer cells (FPC). **(G)** 3D diffusion map and principal curve analysis of cells in (F), showing that FPC and TPCS branched early in the development of BsP towards Bs2/3. **(H)** Cell cycle score of each cell population in (F). BsP, FPC and TPCS are major proliferating populations. **(I)** Marker gene expression for cell types in (F). TM4SF1 marks both BsP and TPCS. Whilst HES1/4 are high in all basal cell lineage, they are both lost in TPCS. In contrast, TPCS specifically upregulates multiple keratins (KRT5/6A/6B/14/16).

The bifurcation of *HES1/HES4* expression levels in basal and superficial (intermediate/umbrella) cells suggests that Notch signalling might contribute to urothelium cell lineage specification (Fig. 2E). In healthy bladder tissue, the expression of Notch ligand *JAG1* was high in endothelial cells and fibroblasts compared to the epithelial cells. Meanwhile, basal cells expressed more *NOTCH3* compared to superficial cells (Fig. 2E). Cell-cell signalling between the urothelium epithelial cells suggested that whilst only Bs1 provides EGF signal, and only intermediate cells provide TGFβ and BMP signal to other cells, Notch signalling is mutual between basal and superficial lineage cells (Fig. 2F). In concordance to previous reports that fibroblasts communicate with BLCA^24, 29^, these results suggest that basal/superficial cell lineage bifurcation might be driven by Notch signalling. The physical proximity of basal cells to the underlying endothelial and fibroblast cells, which provides JAG1/DLL1 ligand, might determine their fate (Fig. 2G).

### Transformation of bladder cancer cells from normal epithelium

Strong transcriptional heterogeneity of BLCA epithelial cells masked underlying common molecular determinants of their evolution. To study BLCA evolution from normal urothelium epithelial cells, genome-wide DNA methylation and single cell chromatin accessibility (scATAC) sequencing were performed (Methods). After quality control, 376,740 cells were obtained in scATAC data^30^, including 107,380 epithelial cells (Supplementary Table 4 and 6-7). To avoid influence from CNV, we clustered the non-immune cells only with variable features from copy-number stable regions (Methods). To this end, 15 clusters of single cells with distinct chromatin accessibility landscapes on differentially methylated regions were identified and termed as ‘epigenotype’ (Supplementary Fig. 3B, Supplementary Table 2 and 9). The distribution of epigenotypes in each tumour was associated with BLCA clinical stages (Supplementary Fig. 3C). Epigenotype Cluster1-4 were stromal cells, whilst Cluster5-15 were epithelial cells (Supplementary Fig. 3B and D). By integrating scRNA-seq data with scATAC, RNA counterparts were matched with epigenotypes (RNA clusters in Supplementary Table 6): normal urothelium basal cells correspond to Cluster14; RNA-determined TPCS cells correspond to Cluster5-7; basal-like cancer cells correspond to Cluster8-13 and luminal-like cancer cells correspond to Cluster15. Similar to the scRNA-inferred developmental trajectory of cancer cells, pseudotime tracing showed that all cancer epithelial cells developed from normal-like Cluster14 cells in scATAC (Methods and Supplementary Fig. 3A).

It is well known that oncogenesis is accompanied by a drastic change in DNA methylation landscape. We found that genomic regions that are hypermethylated in BLCA (hyperDMR) were generally open in stromal cells and remained closed in all epithelial cells regardless of their malignancy states; in contrast, genomic regions that are hypomethylated in BLCA (hypoDMR) is only open in cancer but not normal epithelial cells (Fig. 3A-C and Supplementary Table 2). As DNA methylation is usually accompanied by an inaccessible chromatin state, these results suggest that DNA methylation levels on most hyperDMR are invariable during oncogenesis and the apparent hypermethylation of these regions in tumour tissue is a mere result of cancer clonal expansion. In contrast, active DNA demethylation occurs on hypoDMR accompanied oncogenesis. Hence, hyperDMR can be used as a tool for lineage tracing.

We identified a core subset of hyperDMR which can distinguish MIBC and NMIBC by unsupervised hierarchical clustering. NMIBC Ta and T1-T4 tumours could be classified by DNA methylation levels on different groups of core hyperDMR (Fig. 3D), suggesting they might have distinct cell-of-origins. By correlating gene expression and DMR methylation, we find that NMIBC-specific-hypermethylated regions are associated with basal-specific, superficial-low genes (*GPX4/PRKCZ/CD44*), suggesting that NMIBC originates from superficial cell lineage. On the contrary, MIBC-specific-hypermethylated regions are associated with genes with lower expression in basal cells. Interestingly, a subgroup of T3/T4 MIBC is hypermethylated in all core hyperDMR and correlates with BsP origin by associated gene expression, indicating BsP originated cancer cells are most malignant (Fig. 3D-E). Taken together, these results suggest a heterogeneous origin of BLCA: NMIBC originates from superficial urothelium cells and MIBC develop from basal cells. Finally, the developmental trajectory of MIBC evolution (Fig. 3F) showed that TPSC and FPC are both bifurcated from basal cancer cells (Ca) (Fig. 3G). Whilst most differential expressed genes in FPC are proliferation-related, downregulated Notch signalling and upregulated keratin genes expression in TPCS suggested a phenotype shift (Fig. 3H-I).

Cell-cell signalling analysis showed that FPC specifically received VEGF signals from Ca, and TPCS specifically received BMP signals (BMP2/4 and GDF5) from Ca (Fig. 4A), suggesting BMP signalling might be the extrinsic driver for TPCS genesis. Non-negative matrix factorization (NMF)^31, 32^ identified three independent gene sets (metagenes) NMF6, NMF11 and NMF16, which are highly expressed in TPCS compared to Ca and FPC (Fig. 4B-C). Promoters of these metagenes were enriched with transcription factor binding sites (TFBS)^33, 34^ of SMAD4, ASCL2, NFIB, MEF2C, GLI1, BACH1, BACH2 and BATF (Supplementary Fig. 4-5), suggesting that they are co-regulated (Supplementary Table 7). While NMF16 genes are under the control of NFIB, GLI1/3 and MEF2C, NMF6 and NMF11 genes are directly controlled by BMP via SMAD4 and Notch via ASCL2 (Fig. 4D). As *ASCL1/2* expression is difficult to be assessed in scRNA-seq, we used scATAC data to infer their transcriptional activity. In concordance with the NMF result, ASCL1/2 and SMAD4 TF activities are increased in TPCS without apparent transcriptional upregulation (Fig. 4E).

**Figure 4.**
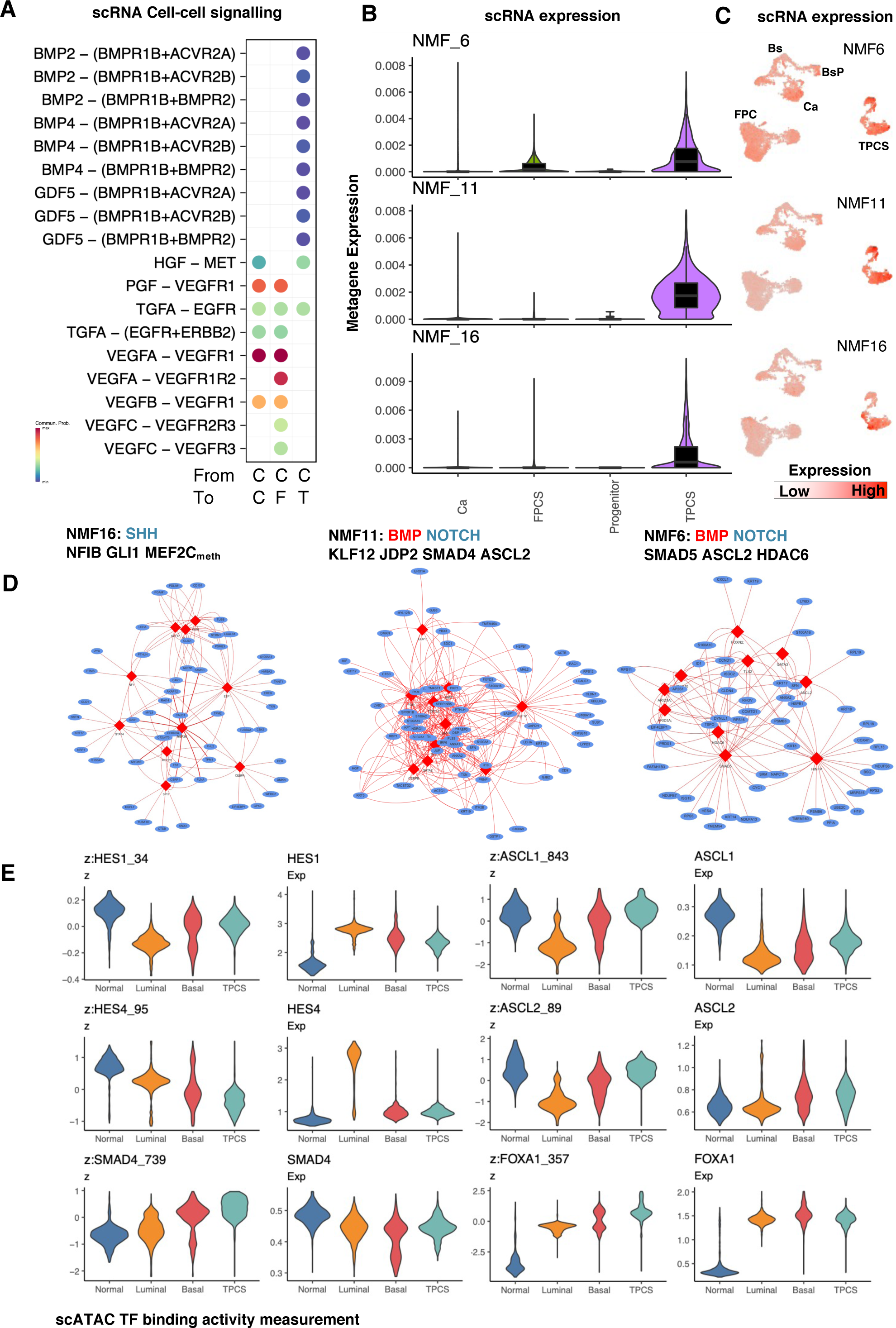
Extrinsic and intrinsic determinants in TPCS transformation. **(A)** Cell-cell signalling probability between BLCA cells in (Fig. 2F). C: Cancer; F: FPC; T: TPCS. BMP signalling distinguished TPCS from cancer and FPC. **(B-C)** Non-negative matrix factorization (NMF) component (“metagene”) expression in single cells. NMF components 6, 11 and 16 are selectively upregulated in TPCS. **(D)** Co-regulated transcription networks extracted from the NMF component by common transcription factor binding site in their promoters showing extracellular signalling event controlling them. Both NMF6 and 11 are controlled by BMP and Notch (co-regulated by SMAD4/SMAD5/ASCL2). NMF16 is controlled by SHH and intrinsically regulated by DNA methylation (co-regulated by NFIB/GLI1/MEF2C). For details of the putative regulatory TF-gene relationship, please refer to Supplementary Table 7. **(E)** TF binding activity (z) and expression (Exp) in normal, luminal cancer, basal cancer, and TPCS from scATAC data.

**Figure 5.**
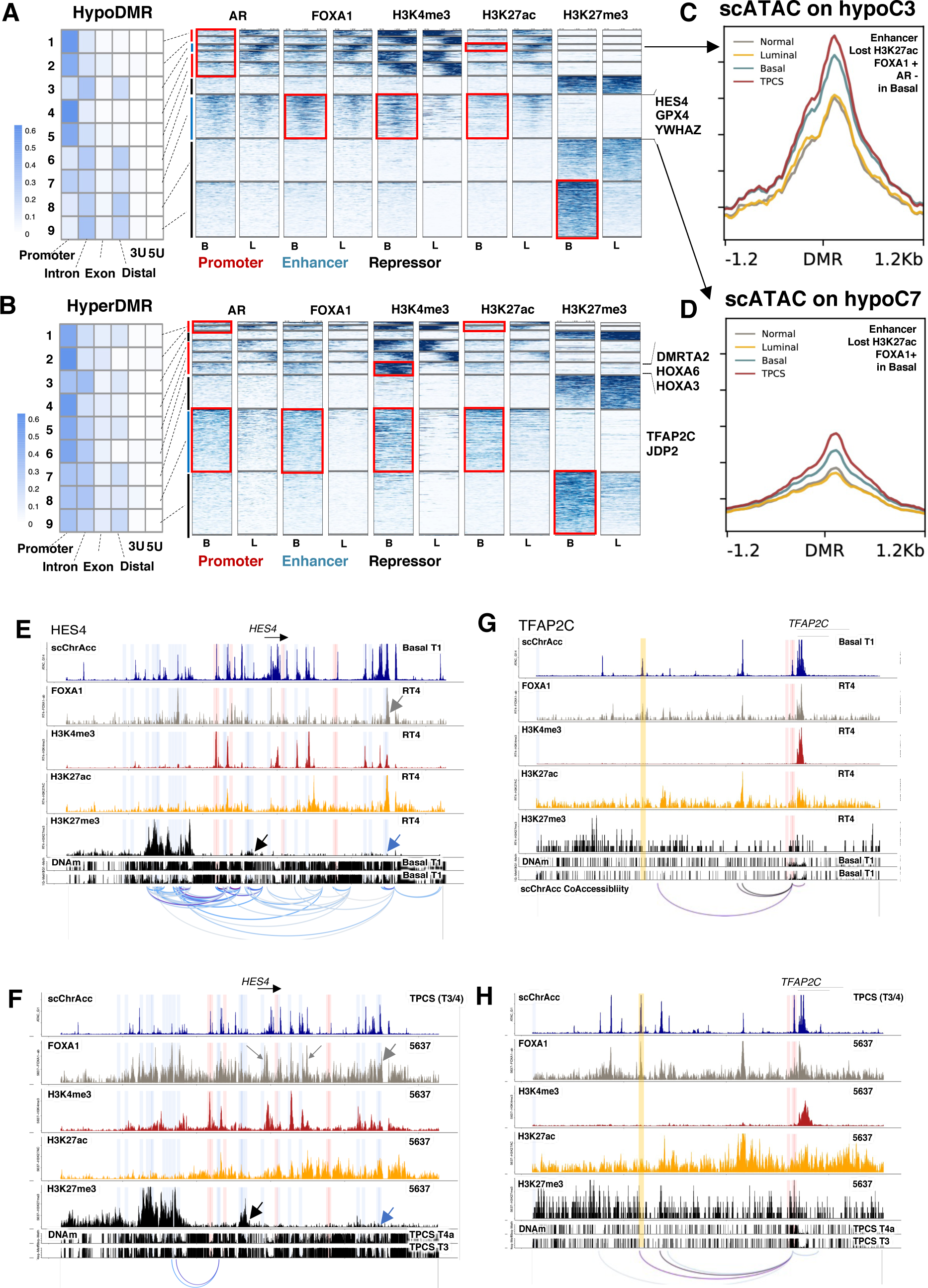
Epigenetic reprogramming in TPCS transformation. **(A-B)** Cut&Tag profiling on MIBC-hypo DMR (A) and MIBC-hyper DMR (B). Left: distribution of DMR clusters on regulatory genomic elements. DMR were K-means clustered by epigenetic modification. Right: Cut&Tag profile of histone modification (H3K4me3, H3K27ac and H3K27me3) and AR/FOXA1 binding in each DMR clusters. Bar annotation on the left denotes regulatory elements (red, promoter; blue, enhancer and black, repressor) **(C-D)** scATAC coverage of hypoC3 (C) and hypoC7 (D). DMR in hypoC3 bound by FOXA1 in both basal and luminal cells but lost H3K27ac modification and AR binding in basal cells (red box in B). DMR in hypoC7 bound by FOXA1 and lost H3K27ac in basal cell. **(E-F)** Epigenetic landscape of HES4 in basal T1 cells and RT4 (E) as well as TPCS and 5637 cell (F). **(G-H)** Epigenetic landscape of TFAP2C in basal T1 cells and RT4 (G) as well as TPCS and 5637 cell (H). Epigenetic features in F-I includes: chromatin accessibility (blue), FOXA1 binding (grey), histone modification (H3K4me3, red; H3K27ac, orange; H3K27me3, black) and DNA methylation (beta), DNAm.

### Emergence of a transcriptional plastic cancer subpopulation by epigenetic reprogramming

We investigated genome-wide distribution of the MIBC master transcription factors FOXA1^35^ and AR as well as H3K4me3, H3K27ac and H3K27me3 on BLCA cell lines (Fig. 5A-B). Two cell lines were chosen for Cut&Tag sequencing analysis: RT4 cell line with *TACC3-FGFR3* representing NMIBC (Luminal, L), and 5637 cell line representing MIBC (Basal, B) (Fig. 5A-B and Supplementary Table 8). Of particular note, as 5637 expressed a high level of *KRT6A/B*, we consider it is a basal-like cell line that mimics TPCS.

Histone modifications and transcription factor binding profiles grouped MIBC-specific hypoDMR into 10 clusters (Methods). MIBC-hypoDMR group3 (hypoC3) lost AR binding as well as H3K27ac modification, and the chromatin accessibility on hypoC3 is increased in basal cancer cells compared to luminal cancer cells, and further increased in TPCS (Fig. 5C). MIBC-hypoDMR group7 (hypoC7) is also identified as enhancers that are bound by FOXA1 and lost H3K27ac in basal cells (Fig. 5D). This group of enhancer controls several basal-low genes such as *GPX4* and *YWHAZ*, as well as *HES4*. Chromatin accessibility on hypoC7 and hypoC3 is highest in TPCS (Fig. 5C-D). On the other hand, MIBC-hyperDMR group8 (hyperC8) is a group of FOXA1-bound, H3K27ac/H3K4me3-high enhancers (Fig. 5B), which controls the TPCS-specific transcription factor JDP2 and TFAP2C; and MIBC-hyperDMR group6 (hyperC6) is a group of promoters that is only active in basal cancer cells that control lineage specification genes such as *HOXA3/HOXA6/DMRTA2* (Fig. 5B). Furthermore, additional loci that were differentially methylated in MIBC are shadowed by H3K27me3 in basal cells only, suggesting PRC2-dependent regulation. In concordance with this result, we observed YY1 binding activity is significantly downregulated in TPCS (Supplementary Fig. 4).

Such epigenomic landscape shift has an immediate effect on gene expression. In TPCS-enriched T3 and T4a samples, DNA demethylation around a distal FOXA1 and CTCF-binding loci 3’ to HES4 lead to loss of long-range chromatin interaction, loss of enhancer activity, and gain of repressive H3K27me3 mark around *HES4* locus (Fig. 5E-F). Similarly, differential FOXA1 binding and DNA methylation led to increased interaction between FOXA1-bound active enhancer to *TFAP2C* promoter in TPCS (Fig. 5G-H). Together, these results suggest that histone modification and DNA methylation changes occur concomitantly with differential AR/FOXA1 binding sites, and might control TPCS-specific transcription programs.

As clinical stage advanced, the predominant population of epigenotype shifted towards TPCS (Fig. 6A). Matching TPCS cells to their CNV profile and integrated RNA data showed that dormant normal cells, proliferating normal cells, and fast-proliferating cancer cells have distinct epigenotype profiles. Whilst the majority of TPCS cells from scRNA clustered to one single CNV type, possibly due to sampling bias, epigenotype of these TPCS cells could correspond to multiple different clusters of scRNA cells (Fig. 6B). In general, the diversity of RNA expression matched to TPCS epigenotype is higher than those matched to other basal cancer cell epigenotypes (Fig. 6C). Orthogonal validation using gene expression diversity analysis (CytoTRACE) showed that FPC and TPCS are likely to be the most undifferentiated, stem-like cells compared to basal and luminal cancer cells (Fig. 6D). Overall, these results suggest that the epigenomic reprogramming in TPCS created a fluid epigenotype that supports multiple transcriptionally diverse phenotypes. In other words, TPCS epigenotype creates ITH without further genetic mutations.

**Figure 6.**
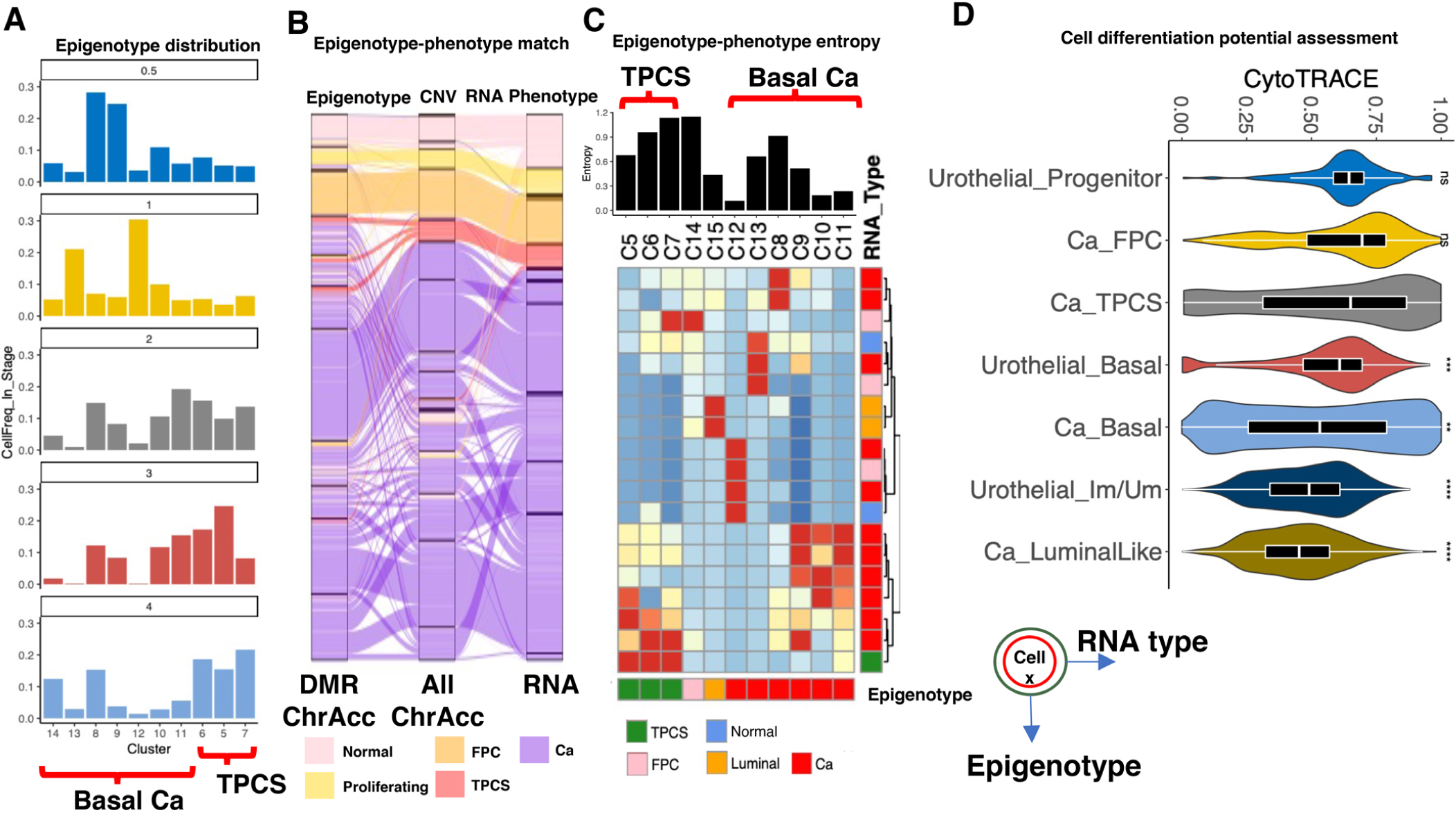
Epigenotype of TPCS is phenotypically plastic. **(A)** Prevalence of ‘epigenotype’ across T stage of BLCA. Epigenotype clusters are clustered by chromatin accessibility on MIBC DMR. Cluster8/9/10/11/12/13/14 are basal cancer and Cluster5/6/7 are TPCS. **(B)** Flow map from epigenotype (left) to copy number variance (CNV, middle) and RNA phenotype in integrated scATAC data. Cells are coloured by their RNA phenotype (bottom). **(C)** Lower: heatmap of single cell epigenotype (x, columns) matching to single cell RNA cluster (y, rows). RNA TPCS cells matched only to epigenotype cluster 5/6/7, which are labelled as TPCS-associated epigenotype. Top: entropy in each epigenotype cluster shows that TPCS-associated epigenotypes match significantly more RNA phenotype than non-TPCS-associated epigenotypes. **(D)** Cytotrace-inferred differentiation potential (high: 1, low: 0) of each epithelial cell types. Differences between groups were measured by adjusted Wilcox test (***: P<0.001, **: P<0.01, *: P<0.05). Compared to TPCS, FPC and the urothelial progenitor cells are stem-like, and basal and luminal cancer cells are more differentiated.

### TPCS heterogeneity supports immune evasion and chemotherapy resistance

ITH created by TPCS might be beneficial for the survival of cancer cells. On one hand, ITH might facilitate immune evasion by shifting neoantigen expression or directly upregulating immune suppression molecules. On the other hand, ITH might facilitate drug resistance by creating a pool of diverse cancer cells that responses differentially to therapy. We studied the immunoediting in tumour tissues by characterizing intratumoral T-lymphocyte population and neoantigen expression in cancer cells. T-lymphocyte clonal expansion status was defined by TCR clone sequence prevalence across tissues. Clonal expansion of T-lymphocyte was absent in healthy or NMIBC Ta BLCA donors but present in T1-T4 donors, and prevalence of clonally-expanded T-lymphocyte and diversity of TCR clone correlates with T-stage, suggesting that whilst bystander (bs) T-lymphocyte infiltration is prevalent in BLCA, they do not mask real tumour-reactive (tr) T-lymphocyte responses, as clonal expansion is still robust (Fig. 7A-B).

**Figure 7.**
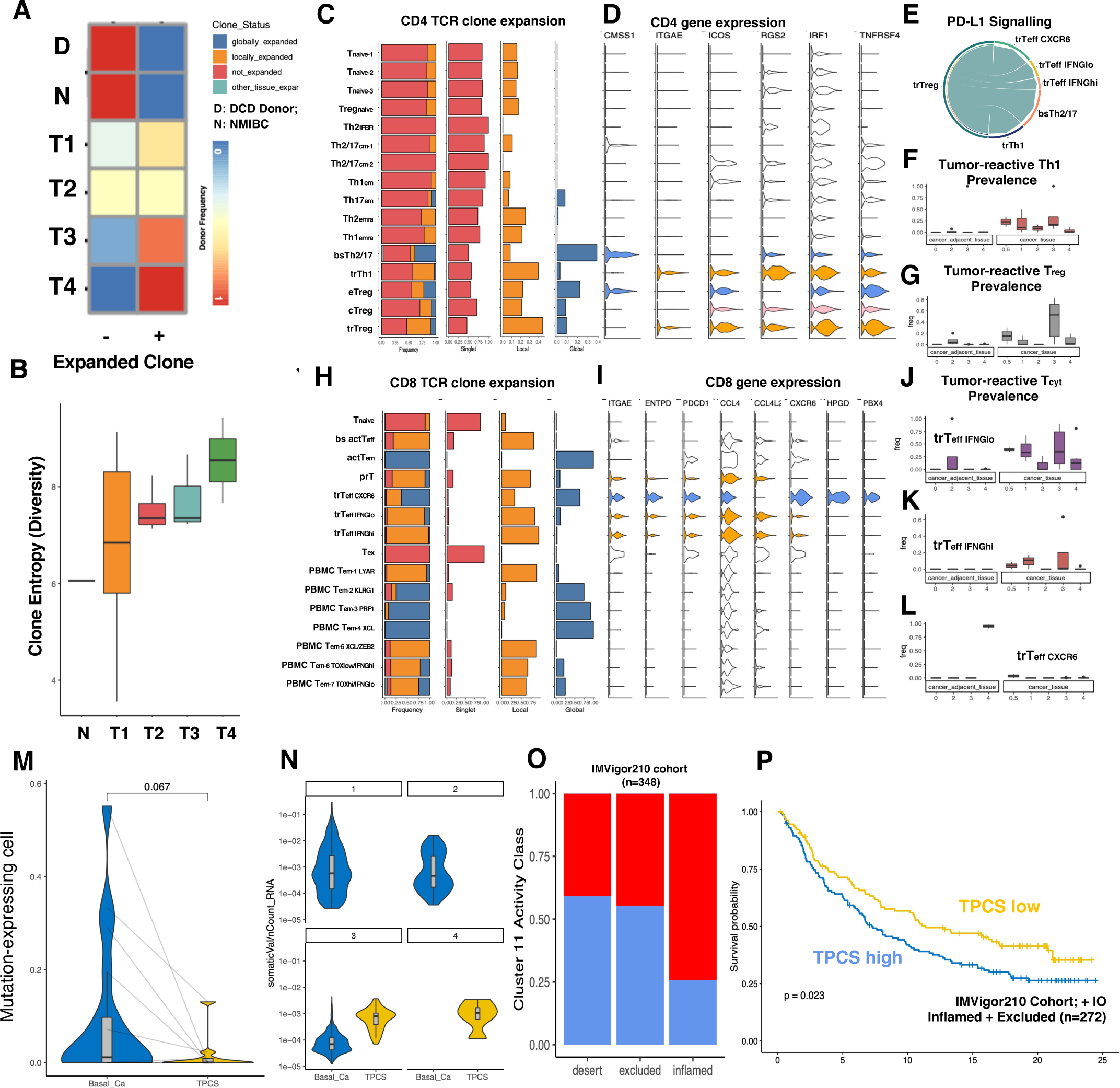
TPCS dynamically regulates neoantigen expression to evade immunoediting. **(A)** The prevalence of T cells with(+)/without(-) clonal expansion across BLCA T stages. 100% advanced T stage BLCA patients have clonally expanded T cells in tumour tissue. **(B)** TCR clonotype diversity (Shannon entropy) across clinical stages shows a general trend of increase in clonotype diversity in the advanced clinical stage. **(C-D)** Clonal expansion (C) and gene expression (D) of each class of CD4 T helper cells. Colour in (C): red, singlet; orange, local-expanded (>2 similar TCR found only in bladder tissue); and blue, global-expanded (>2 similar TCR found in multiple tissue samples from the same donor). Colour in (D): blue, global-expanded; pink and orange, local-expanded. **(E)** PD-L1 cell-cell signalling with all bladder tissue cells revealing that trT_reg_ is the main, if not the only one, contributor of PD-L1 ligand for other tumour-reactive T cells. **(F-G)** The prevalence of trTh1 (F) and trT_reg_ (G) in cancer and adjacent tissues. **(H-I)** Clonal expansion (H) and gene expression (I) of each class of CD8 cytotoxic T cells. Colour annotation is the same with (C-D). **(J-L)** The prevalence of trT_effIFNGlo_ (J), trT_effIFNGhi_ (K), and trT_effCXCR6_ (L) in cancer and adjacent tissues. **(M)** Frequency of single cancer and TPCS cells expressing mutation-carrying transcripts in each donor. **(N)** Normalized per-cell mutation-carrying transcript expression level, separated by clinical stage. **(O-P)** TPCS metagene activity (O) and survival probability (P) in IMVigor210 cohort. Samples are classified as immune desert, excluded, or inflamed as in the original paper.

For CD4^+^ T-cell, clonal expansion is more prevalent in the mature T-cell population (bsTh2/17, trTh1, cT_reg_, eT_reg_, trT_reg_) (Fig. 7C and Supplementary Fig. 6). Among these CD4^+^ T-cell, globally-expanded population (bsTh2/17, eT_reg_) do not express *ITGAE*(CD103) and *ENTPD1*(CD39) (Fig. 7D and Supplementary Fig. 6). On the contrary, locally-expanded trTh1 and trT_reg_ overexpress *ITGAE/ENTPD1* in addition to *RGS2/IRF1/TNFRSF4* (Fig. 7D and Supplementary Fig. 6). In the light of previous reports^36^ and considering the local TCR clone expansion, these results suggested that CD103/CD39 combination might serve as a general marker for tumour-reactiveness^37^. Cell-cell signalling analysis suggests that trT_reg_ contributed a majority of PD-L1 ligand to other T cells in the tumour environment (Fig. 7E). The prevalence of trT_reg_ increased in T3 whilst trTh1 decreased over clinical stage progression (Fig. 7F-G). For cytotoxic CD8^+^T cells in tumour environment, we found that the globally-expanded cells include the immature, non-PD-1-expressing T_emra_(KLRG1), T_emra_(PRF1), T_emra_(XCL) and PD-1 expressing (non-locally-expanded) actT_eff_(GZMK) (Fig. 7H-I and Supplementary Fig. 7). Clonal expansion together with PDCD1 expression classified four clusters of T_eff_: the *ZEB2^+^/TOP2A^+^* proliferating prT_eff_, IFN_Ghi_trT_eff_ and IFN_Glow_trT_eff_, and *IFNG^-^/CXCR6^+^* trT_eff_ (trT_eff_ CXCR6) (Fig. 7H and Supplementary Fig. 7). These trT_eff_ were more prevalent in cancer tissues, with trT_eff_ CXCR6 is the only *PDCD1^+^/ENTPD1^+^/ITGAE^+^* population that contains globally-expanded clones and might support strong antitumor activity^38^ (Fig. 7H, J-L and Supplementary Fig. 7).

The above results suggest BLCA tumour microenvironment is not completely immunosuppressive and continues to attract and train new naïve T cells to tumour-reactive controllers. Hence, BLCA might evade immunosurveillance via alternative mechanisms. We analyzed MHC-related molecule expression and mutation-carrying (‘neoantigen’) transcript expression in cancer cell scRNA data (Supplementary Table 9). Whilst MHC-related molecule expression is higher in TPCS than basal cancer cells, marginally significant fewer TPCS expressed mutation-carrying transcript than basal cancer cells (Fig. 7M). In basal cancer cells, mutation-carrying transcript expression level reversely correlates with the clinical stage. In contrast, TPCS expressing mutation-carrying transcripts remained stable (Fig. 7N). These results suggest that TPCS switch between mutation-high and mutation-free states to evade immunosurveillance. Finally, the gene-set activity of TPCS marker metagene NMF11 component was used to quantify TPCS prevalence in BLCA from IMvigor210 dataset. Compared to immune-desert tumours, inflamed tumours are more likely to contain more TPCS (Fig. 7O). In the group of patients who received immunocheckpoint inhibitor (ICI) treatment, patients with high TPCS marker gene expression have significantly poorer prognosis (Fig. 7P), suggesting that TPCS contributed to ICI resistance.

A subgroup of patients in our cohort received adjuvant intravesical gemcitabine instillation. Whilst epigenotype-related phenotype diversity does not change between treatment-naïve and post-chemotherapy patients for TPCS (Fig. 8A), the prevalence of TPCS but no other basal cancer cells is significantly increased after chemotherapy in different clinical stages (Fig. 8B-D). Cell cycle score revealed that after gemcitabine treatment, the majority of basal-like and luminal-like cancer cells tend to be arrested in G1 and do not enter S or G2/M phases (Fig. 8E). Additionally, FPC tend to be arrested in G2/M, and TPCS become more active as the prevalence of both S and G2/M-phase cells increased (Fig. 8E). In TCGA BLCA cohort, where patients were mainly treated with chemotherapy, high TPCS marker gene expression significantly correlates with poor prognosis (Fig. 8F), suggesting that TPCS contributed to chemotherapy resistance.

**Figure 8.**
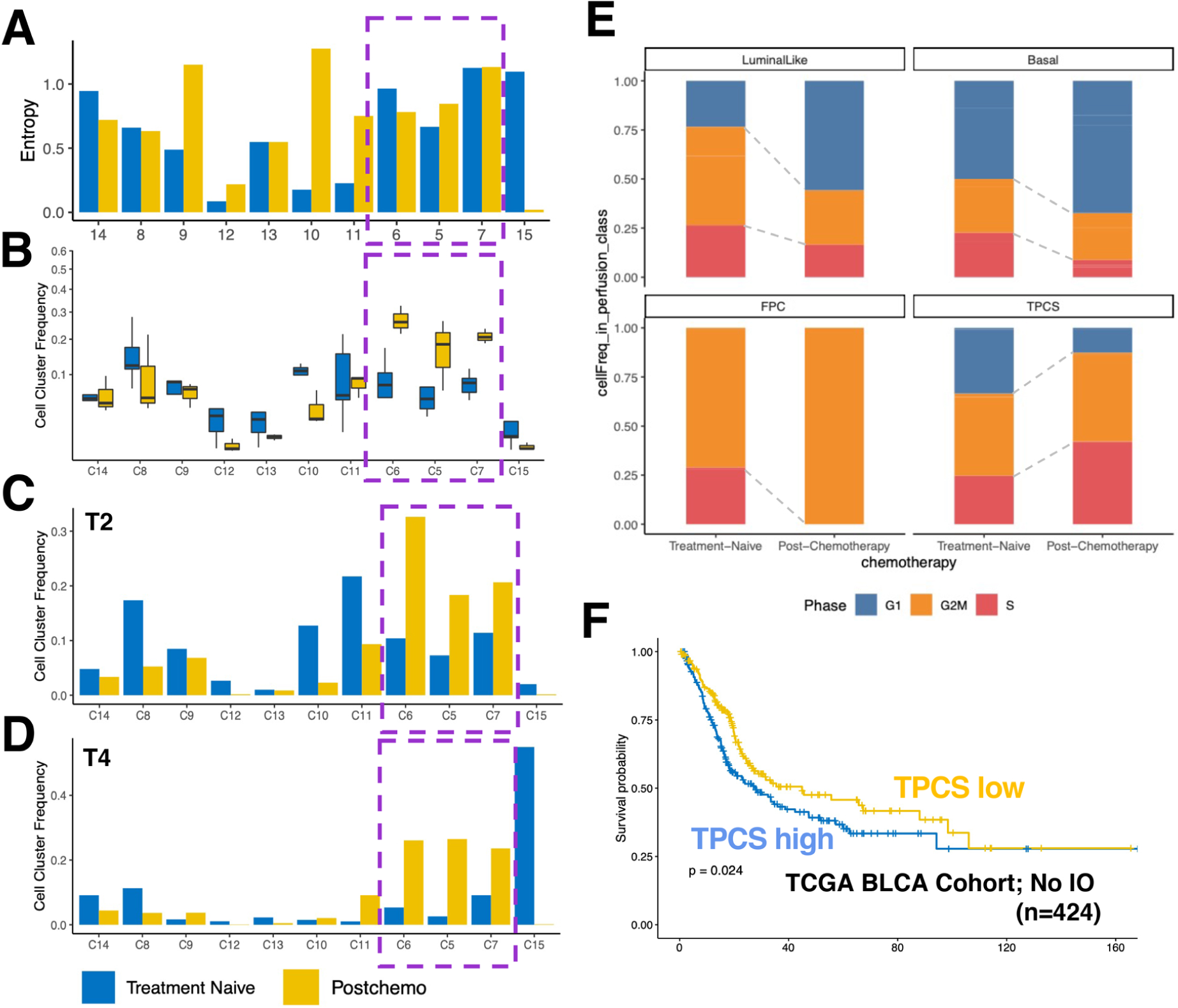
TPCS dynamically shifts cell cycle state to escape from chemotherapy. **(A)** Epigenotype-phenotype matching entropy for each epigenotype cluster. Cells are separated by their donor into treatment-naïve and post-chemotherapy groups. TPCS has high epigenotype plasticity regardless of chemotherapy induction. **(B)** Epigenotype prevalence in treatment-naïve and post-chemotherapy donors, showing that TPCS epigenotype is increased after chemotherapy treatment. **(C)** Epigenotype prevalence in stage 2 patients. **(D)** Epigenotype prevalence in stage 4 patients. **(E)** Cell cycle phase of basal, luminal-like, FPC and TPCS cancer cells in treatment-naïve and post-chemotherapy samples. TPCS shows lower proliferating potential in the treatment-naïve stage than FPC, but is the only cell type with elevated proliferating potential post-chemotherapy. **(F)** Survival probability of patients with high and low TPCS metagene activity in TCGA bladder cancer patient cohort

## Discussion

BLCA is a complicated and heterogeneous disease. The evolution of BLCA from normal urothelium to generate such heterogeneity remains unclear^20^. Particularly, no direct evidence was made from the molecular level to clarify the origins of basal/luminal types of human BLCA. Through DNA methylation based lineage tracing and scATAC-seq, our results revealed a comprehensive picture of BLCA evolution. NMIBC and MIBC originate from superficial intermediate/umbrella or basal cells of the urothelium, respectively. Cell-of-origin of BLCA corresponds to their clinical behaviour, with BLCA originating directly from BsP being most malignant. In late stage, a progenitor-like, *TM4SF1*-positive cancer cell subpopulation emerge from the MIBC track. TPCS is induced by epigenetic reprogramming, and contributes to ITH by supporting highly plastic transcriptional phenotypes (Supplementary Fig. 8).

TM4SF1, a member of the tetraspanin family, has been identified as a tumour-specific antigen, promoting proliferation, invasion, epithelial-mesenchymal transition (EMT) and chemo-resistance^39–41^. Expression of TM4SF1 marks stem cell in mesenchymal tissue^42^ and lung^43^. Here, we identified that TM4SF1 also marks human bladder urothelium progenitor cell. Additionally, TM4SF1-positive cancer cell subpopulation is also highly proliferative and transcriptionally plastic. TPCS is phenotypically similar to hyper-plastic cancer cells reported in other cancer types^7, 8, 44, 45^. In our dataset, all metastatic, squamous differentiation BLCA-isolated cells are TPCS. TPCS expresses lower HES1, HES4 and Notch pathway downstream targets, possessing the characters of cancer stem-like cells (CSCs). Consistent with the notion that the phenotype of CSCs is often regulated by epigenetic modifications^44^, our findings suggested that epigenomic reprogramming in TPCS results in a variety of distinct transcriptional phenotypes. These data suggested that during BLCA progression, the adaptation of a progenitor/stem-cell like fate occurred and contributed to ITH in BLCA. The emergence of TPCS increased phenotypic diversity in tumours and facilitated immune evasion and chemotherapy resistance (Supplementary Fig. 8). Our study provides not only insights into BLCA biology but also novel clinical targets for BLCA therapy.

## Supporting information

Supp.Table.1. Human biospecimen and experiment details

Supp.Table.2. MIBC associated DMR loci

Supp.Table.3. scRNA marker genes

Supp.Table.4. scATAC marker peaks

Supp.Table.5. Cell Annotations for scRNA.

Supp.Table.6. Cell Annotations for scATAC epithelial cells

Supp.Table.7. cisTarget-inferred TF-to-gene matches for NMF co-regulatory gene network in TPCS

Supp.Table.8. Cut&Tag peaks in RT4 and 5637 cells

Supp.Table.9. CNV of sequenced tumours

Supp.Table.10. SNV of sequenced tumours

## Methods

### Human biospecimens and cell lines

This study was conducted in accordance with the measures of China on the administration of clinical research, the Declaration of Helsinki, and the Institutional Ethic Protocols of Zhongnan Hospital of Wuhan University (ZHWU), Beijing Friendship Hospital, Peking University International Hospital and Hubei Cancer Hospital. The study protocol was approved by the Institutional Review Board (IRB) at ZHWU (approval number: 2015029). All patients and all relatives of organ donors provided written informed consent. All study procedures were performed in accordance with the ethical standards of the Institutional Ethics Review Committee. Human sample preservation by the Department of Biological Repositories at ZHWU, the official member of the International Society for Biological and Environmental Repositories (https://irlocator.isber.org/details/60), was approved by the IRB (approval number: 2017038) at ZHWU and China Human Genetic Resources Management Office, Ministry of Science and Technology of China (approval number: 20171793).

Clinical, pathological, and follow-up data records for all patients were collected (Supplementary Table 1) and three pathologists were invited to independently confirm the histology diagnosis. Additional cancer patient samples were collected through the clinical practice of the ZHWU, Beijing Friendship Hospital, Peking University International Hospital and Hubei Cancer Hospital. The study uses of clinical information and human samples (including blood, surgical tissue specimens, and primary cancer cells) was approved by the Institutional Ethics Review Committee (approval number: 2015029). Routine laboratory tests and pathology assessments were done according to the relevant Chinese clinical protocols.

RT4 and 5637 cells were kindly provided by Cell Bank, Chinese Academy of Sciences (Shanghai, China) and cultured under identical conditions following standard procedures^46^. To construct 5637-AR and RT4-AR stable cell lines, the LV6-AR lentivirus packaging was provided by GenePharma Inc (Shanghai, China). The lentivirus particles were used for infection of target cells with the supplement polybrene. After two rounds of infection, cells were selected with 1 μg/ml puromycin. Clinical assessment of bladder cancer was done according to the EAU 2020 Oncology Guidelines (https://uroweb.org/individual-guidelines/oncology-guidelines/). Human peripheral blood was collected with BD EDTA tube according to the manufacturer’s protocol and stored at 4 degree Celsius for no longer than 8 hours before serum separation. Peripheral blood mononucleus cell (PBMC) was separated with the standard Ficoll protocol. Fresh tumour or normal tissues was collected during surgery and transferred to laboratory in high glucose, 10% FBS supplemented DMEM. Tissue sample was resected in PBS prior to single cell dissociation. For methylation sequencing, tissue samples were flash frozen in liquid nitrogen and stored at -80 degree Celsius.

### Public data

The study used a series of human bladder cancer sequencing data from NCBI SRA sequencing read archive (https://www.ncbi.nlm.nih.gov/sra) with the accession code PRJNA662018. TCGA BLCA dataset from UCSC XENA (http://xena.ucsc.edu) were used in the study. The IMVigor210 dataset were downloaded from http://research-pub.gene.com/IMvigor210CoreBiologies/packageVersions/. The Encode transcription factor binding sites, ChromHMM tracks, CpG islands, repeat regions data, and lift-over chains were collected from UCSC Genome Browser (http://www.genome.ucsc.edu/). Transcription factor binding information were downloaded from ReMap database (http://pedagogix-tagc.univ-mrs.fr/remap).

### Statistical methods

No statistical methods were used to determine the sample size used in the study. Between-group statistics was done with T-test (two-sided) or Wilcoxon Rank Test if T-test was not applicable. Survival analysis was done by Cox method. Detailed statistical methods were briefly denoted in the figure legends or text accompanying. All statistical analysis in this study were performed using R (3.6.2) (http://CRAN.R-project.org).

### Molecular Biology

#### Nucleic acid preparation

For methylation sequencing or mutation panel capture sequencing, genomic DNA was extracted from tumour or normal tissue with Qiagen Animal Tissue DNA Extraction Kit (Qiagen Cat. #69504) according to the manufacturer’s protocol. Genomic DNA from FFPE tissue slides were extracted using MagPure Tissue DNA DF Kit (Magen Inc., Cat. #MD5112-TL-06). Extracted DNA were quality-controlled by Qubit dsDNA HS assay (Thermo Fisher Scientific) and Agilent 2100 Fragment Analyzer.

#### Chromatin accessibility sequencing (ATAC-seq) on tumour tissue

20mg flash-frozen tumour tissue samples were minced using a double-sized douncer (Sigma, Cat. #D8938) in 1xHB (0.25M sucrose, 0.06M KCl, 0.015M NaCl, 0.005M MgCl2, 0.01M Tris-HCl pH 7.5), added to 5ml trypsin and 40µl 5U/µl DNase I (Sigma, Cat. #D5025) and digested in 37 degree Celsius for 45 mins, with two times of rotation in between to mix the reaction. The digested cells were then neutralized with equal volume DMEM (Thermo Fisher, Cat. #11995065) plus 10% FBS (Gibco, Cat. #16000044) and filtered through a 70µm cell filter (BD Falcon, Cat. #352350). The homogenize was centrifuged at 500x g, 4 degree Celsius for 5 mins. The sedimented cells were then resuspended in 400µl 1xHB and washed once, transferred to 2ml LoBind Tube (Eppendorf) and washed again. Cells were counted using Trypan blue (Solarbio, Beijing, China). After quantification, the cells were then added to a 30%-40%-50% iodixanol (Sigma, Cat. #D1556) gradient and centrifuged at 3000 g, 20 mins at 4 degree Celsius. The cell layer at 30%-40% interface was collected for library preparation. DNA library were prepared (‘tagmentation’) with a Tn5 transposase kit (Vazyme, Cat. #TD501) using 1 million cells per reaction according to manufacturer’s protocol. After tagmentation and PCR amplification, the sequencing library were quality-controlled with SYBR-green based qPCR using primers for house-keeping gene (GAPDH) promoter and gene desert (chr5: 105187030-105190000) before sequencing.

#### Cut&Tag sequencing on cells

RT4 or 5637 cells were counted using Trypan blue (Solarbio, Beijing, China). After quantification, 40 million cells were used for Cut&Tag experiment. Cut&Tag experiment were performed with NovoProtein Cut&Tag 2.0 pAG-Tn5 kit (NovoProtein, Cat. #N259) according to the manufacturer’s protocol. Antibodies used in this study include: anti-H3K4me3 (Diagenode, Cat. #C154100003), anti-H3K27me3 (Abcam Cat. #ab6002), anti-H3K27ac (Abcam Cat #ab4729), anti-FOXA1 (Abcam Cat. #ab170933) and anti-AR (Abcam Cat. #ab108341), and Goat-anti-mouse IgG (Sangon, Cat. #D111024) and Goat-anti-rabbit IgG (Sangon, Cat. #D111018). Each library was sequenced to 2x human genome coverage on Novaseq sequencer (Illumina, CA).

#### Single stranded DNA methylation capture sequencing

Tissue genomic DNA (200ng) were bisulfite converted using EZ-DNA Methylation-Gold Kit (Zymo Research, Cat. #D5006) according to the manufacturer’s protocol. After conversion, the DNA were subjected to a single-stranded library preparation protocol Tequila 7N (Euler Technology). In brief, the DNA were end-repaired using Klenow fragment (NEB) and tailed with poly-A homopolymer using terminal deoxynucleotide transferase (Takara), ligated to a poly-T overhang adaptor using T4 DNA ligase (Enzymatics), and linearly amplified for 12 cycles using PhusionU (Thermo Fisher Scientific). The amplified linear products were then annealed to a 5’ adaptor with 7bp 3’ random nucleotide overhang and PCR-amplified using adaptor oligos (Sangon, Shanghai, China) and Phusion (Thermo Fisher Scientific), resulting in a library with proper Illumina sequencing adaptor ends ready for NGS. Hybridization was done with SeqCap EpiGiant Enrichment Probe (Roche, Cat. #07138911001), oligos and SeqCap wash and binding buffers (Roche) following the manufacturer’s protocol. After hybridization, the library was amplified using Phusion for 8 cycles and sequenced on Novaseq sequencer (Illumina, CA) to a target of 100M paired-end 150bp reads.

#### DNA mutation panel sequencing

DNA was sonicated into ∼250 bp fragments with Covaris S220. NGS sequencing libraries were built with a single-stranded DNA ligation protocol. In brief, sonicated DNA was denatured to form a single strand and 3’-polyA-tailing was performed with terminal transferase (Cat. #P7070, Enzymatics Ltd., USA). Ligation of a polyT-extruding adaptor (Sangon Ltd., China) was performed with *E.coli* ligase (Cat. #2161, Takara Ltd., Japan). Linear amplification of the ligated product was performed with adaptor-specific primer (Sangon Ltd., China) for 12 cycles and the amplified product was annealed and ligated into a 5’-polyN-extruding adaptor (Sangon Ltd., China) with T4 ligase (Cat. #L6030, Enzymatics, USA). The ligated product was then amplified with Illumina-compatible primers (Sangon Ltd., China) for 10 cycles. For panel capture sequencing, the amplified library was captured using either: (1) a custom-synthesized oncology panel-consisting exons, UTR and structural variant breakpoint-enriched introns of 538 tumour-related genes, as well as 1076 SNP loci (Euler Technology Ltd., China), or (2) a custom-synthesized whole exon panel consisting exons, UTR and SNP loci from GRCh37 version human genome totalling 52.6Mbp (Euler Technology Ltd., China). Libraries were sequenced to targeting ∼800x (oncology panel) or ∼150x (whole-exome) on-target coverage with paired-end 150bp read format on Illumina Novaseq.

#### Single cell RNA and chromatin accessibility sequencing

Fresh tissues were processed immediately after being obtained from bladder cancer patients. Tissues were cut into tiny pieces (<1mm diameter) and then subjected to dissociation using collagenase II (Biofrox, Cat. #2275MG100) and 100 μl of DNase (Servicebio, Cat. #1121MG010) at 37 degree Celsius for 1hr. After dissociation, cells were filtered with 40 µm BD filter mesh and subsequently centrifuged at 250g x 5min. Cell pellets were washed in PBS twice, and resuspended in 1ml ice cold RBC lysis buffer and incubated at 4 degree Celsius for 10 mins. 10 ml of ice-cold PBS was added to the tube and subsequently centrifuged at 250g x 10min. After decanting the supernatant, the pellet was resuspended in 5 ml of calcium- and magnesium-free PBS containing 0.04% weight/volume BSA. Cells were counted using Trypan blue (Solarbio, Beijing, China). For chromatin accessibility sequencing, approximately 106 cells were used for nucleus extraction.

Nucleus extraction were performed as 10x single cell library preparation was done according to the manufacturer’s protocol. Chrominum Single Cell 3’ V3 kits, 5’ V3 kits (for TCR sequencing), and ATAC V2 kits were used. For RNA sequencing, single-cell suspensions were loaded onto a Chromium Single-Cell Controller Instrument (10× Genomics) to generate single-cell gel beads in emulsions (GEMs) targeting ∼8000 cells (3’/5’ RNA). After generation of GEMs, reverse transcription reactions were engaged to generate barcoded full-length cDNA, which was followed by disruption of emulsions using the recovery agent, and then cDNA clean-up was performed with DynaBeads Myone Silane Beads (Thermo Fisher Scientific). Next, cDNA was amplified by PCR. Subsequently, the amplified cDNA was fragmented, end-repaired, A-tailed, and ligated to an index adaptor, and then the library was amplified. To amplify TCR sequence, 10x Genomic TCR kit was used according to the manufacturer’s protocol. The scRNA libraries were sequenced aiming to have ∼5000 reads per cell on Illumina Novaseq with paired-end 150bp reads.

For ATAC, tagmentation were performed according to the manufacturer’s protocol. After tagmentation reaction, nucleus suspensions were loaded a Chromium Single-Cell Controller Instrument (10× Genomics) targeting ∼10000 nucleus in one reaction. After generation of GEMs, PCR reaction were performed to amplify the library. DNA clean-up was performed with size-selection XP beads. Libraries were sequenced aiming to have ∼5000 reads per cell on Illumina Novaseq with paired-end 50bp reads.

### Bioinformatics

#### Mutation profiling

Raw sequencing data were mapped using bwa-mem to GRCh37 reference genome with default parameters. Germline mutations were called with the Sentieon haplotyper (Sentieon-Genomics-201808.05) and annotated with VEP^47^ (90.1) and SnpSift^48^ (4.2). For paired tumour-normal samples, candidate germline variants were filtered with the gnomAD global frequency <0.001 and in-house database frequency <0.001 (out of 20,000 patients). Intersection with variants found in different male members of the pedigree was performed to extract patient-specific germline mutations. Somatic tumour mutations were called with the Sentieon TNscope (Sentieon-Genomics-201808.05) with paired sample or NA12878 (at unpaired cases) and Pisces^49^ (5.2.9.122), while variants called by both algorithms were passed for filtering. CNV were called using CNVkit^50, 51^ with default parameter. B-allele frequency (BAF) determination was performed for germline and somatic variants. Tumour genome was segmented using BAF and sequencing depth information. Allelic copy numbers were determined for each somatic variant using a hypergeometrical test. Tumour content determined by an in-house CNV-based linear regression method was confirmed by Hematoxylin and Eosin (H&E) staining. Minimal tumour-cell-fraction of 5%/2%, minimal variant read number of 10/3, and minimal read depth of 500/30 were applied to variants for filtering for panel sequencing or whole-exon sequencing, respectively. Filtered mutations were annotated with vcfanno and filtered with gnomAD global frequency <0.001.

#### Genomic region liftover

For genomic regions liftover between UCSC GRCh38 and GRCh37, the R package easyLift (https://github.com/caleblareau/easyLift), the liftover executable from UCSC Kent Utility (https://genome.ucsc.edu/cgi-bin/hgLiftOver) and the lift-over synteny chain files from UCSC Genome Browser were used. Target genome was always GRCh38.

#### DNA methylation data processing

Raw bisulfite-converted DNA methylation sequencing data, either downloaded from NCBI SRA or directly from in-house sequencing, were processed using fastp (--trim-front2 20 -w 20)^52^ (https://github.com/OpenGene/fastp) and mapped to GRCh37+decoy reference genome using BWA-Meth (https://github.com/brentp/bwa-meth) using standard parameters. Mapped data were deduplicated and sorted using Sambamba (https://github.com/biod/sambamba) and Samblaster (https://github.com/GregoryFaust/samblaster). CpG-methylation level was extracted using Pile-O-Meth (https://github.com/dpryan79/MethylDackel) toolkit. For all libraries, conversion rate was quality controlled by CHH methylation level >99%. Basic statistics of in-house sequencing library were further quality-controlled by on-target rate and on-target coverage with bedtools (https://github.com/arq5x/bedtools), and duplication rate and mapping rate with Sambamba.

#### Differential methylated loci and region

CpG methylation level (beta: defined as reads of C nucleotide over total read coverage on single C bases on both strands on CpG loci) was measured for each CpG loci across the genome as mentioned above using Pile-O-Meth. For each loci, beta from sequencing results were summarized in R (3.6.2) using an in-house script. Differentially methylated loci (DML) were defined as: (1) P<0.01 for T-test between control and case groups (given NMIBC/MIBC or BC/normal); (2) beta difference between case and control groups > 0.1. Initial DMR candidate were made by merging within-100bp-apart DML. The average beta of each initial DMR were calculated as mean beta of all CpG encompassed in the DMR. This average beta was subjected to t-test and P<0.01 regions were selected as candidate ‘seed’ DMR. Segments of methylation difference level were computed using a circular binary segmentation approach on beta difference case and control groups with DNAcopy^51^ (https://github.com/veseshan/DNAcopy). K-means clustering was performed using R (3.6.2) on the methylation beta difference on each segment, and clusters of segments fully encompassed candidate ‘seed’ DMR were selected as true DMR candidate.

#### ATAC and Cut&Tag sequencing data preprocessing

Raw paired-end open chromatin tagmentation (ATAC) and Cut&Tag sequencing data were mapped to human reference genome GRCh38 using Bowtie2 (-k 10 --very-sensitive -X 2000) (https://github.com/BenLangmead/bowtie2). All unmapped reads, non-uniquely mapped reads, reads with low mapping quality (MAPQ < 20) and PCR duplicates were removed. For Cut&Tag sequencing libraries, data were used as is. For in-house prepared ATAC-seq data, the data were quality-controlled by assessing insertion size (using an in-house R script) and TSS-enrichment (using an in-house R script with GenomicRanges package (https://github.com/Bioconductor/GenomicRanges) measuring the depth ratio at the promoter region (GRCh38 refFlat annotation from UCSC Genome Browser) (0bp of TSS vs. 1kbp +- of TSS). A QC-passed ATAC-seq library must have TSS enrichment of 6, mapped deduplicated sequencing fragments ≥ 20M PE reads, PCB1>0.9, PCB2>3 (https://www.encodeproject.org/pipelines). Enrichment peaks were determined by intersecting peaks found from MACS2 callpeak (-f BAMPE)^53^ (https://github.com/taoliu/MACS) and Genrich (-r -m 1 -j; for ATAC only; and standard parameter for Cut&Tag) (https://github.com/jsh58/Genrich). Quality control of ATAC-seq libraries including read length, V-plot and TSS-enrichment were done with custom R script and deeptools (https://github.com/deeptools/deepTools). Reliable peaks were identified with IDR (https://www.encodeproject.org/software/idr). Reliable ATAC peaks from different set of data were converged with 1bp minimum overlap and extended to the largest width of overlapping peaks. Joining these operation results in a set of non-overlapping, varied-width peaks across the genome encompassing all reliable open chromatin region.

#### Measurement of difference of ATAC-seq peaks

Read coverage of sequencing library were collected over the repeatable ATAC enrichment peak mentioned above with Sambamba. Differential enrichment was performed with DESeq2^54^ (https://github.com/mikelove/DESeq2) using standard parameters. For samples with few replicates, we adapted a general linear model approach for estimating difference following the method mentioned in Reilly et al 2015^55^.

#### Classification of differentially methylated region by Cut&Tag signal

Reads from Cut&Tag sequencing library were transformed into bigwig using deepTools^56^. Windows +-1200bp around the differentially methylated loci were used to extract the reads. The resulting coverage matrix containing H3K27me3, H3K27ac, H3K4me3, anti-FOXA1 and anti-AR signals were then clustered using K-means with deepTools^56^. Annotation of these K-means clustered regions were done by ChIPseeker^57^ using -1000bp∼+500bp as promoter around TSS, and their relative distribution on different types of genomic regions were plotted using pheatmap (https://cran.r-project.org/web/packages/pheatmap/index.html). Regions were labelled ‘Promoter’ if they were mainly covered by H3K4me3 in any cell type, ‘Enhancer’ if they were covered by H3K27ac in any cell type, and ‘Repressor’ if they were covered by H3K27me3 in any cell type.

Upon these regions we did not detect bivalent promoter/enhancer features with H3K4me3/H3K27me3 double positivity.

#### Single cell RNA preprocessing

Loompy-Kallisto^58^ were used for mapping the RNA data for gene expression analysis. Loom files were read in R by hdf5r (https://cran.r-project.org/web/packages/hdf5r/index.html) and pre- processed with Seurat^26^ 3.0. Quality control was done for every single sample individually to filter against gene counts, UMI counts, total reads, mitochondrial reads. Generally, cells with >10% mitochondrial reads, or with UMI < 600 or > 5000, or with gene counts > 5000 were filtered prior to subsequent analysis. Such quality control process might iterate at every subsequent step to ensure the stringency of analysis. Individual samples were processed through the Seurat pipeline. Data firstly passed DoubletFinder^27^ pipeline with standard parameters to filter against potential doublets. The filtered data were then normalized and top 2000 variable genes were identified. Ribosomal proteins, heat shock proteins, and chaperones were intentionally removed from the variable gene list because of the highly inconsistent nature of their behaviours between different tissue types. Gene expression profile were then scaled and reduced by PCA using Seurat. Generally, ≥30 PCA components were included in subsequent steps. FindClusters function were initially performed using a high resolution then gradually lowered to ensure the final clusters are less than PC components. SingleR annotation with human reference and conventional markers were used to categorize the cell clusters. Differentially expressed genes were identified with Seurat FindAllMarkers function with Wilcoxon test for the cell clusters. Comparison of the found DE genes with conventional markers were performed to ensure that clusters contain relatively pure cell population. Cell-type-annotated cells were then separated into different subsets based on their types. In this study, epithelial cells, endothelial cells, fibroblast, CD8 T cells, CD4 T cells, NK cells, B cells and myeloid cells were considered ‘pools’ for subsequent analysis. After pre- processing, similar type of cells from different samples were merged and re-analyzed. Quality control, doublet identification, dimensionality reduction, cluster identification, differentially expressed gene identification, and cell type identification were iteratively performed on these ‘pure’ sets of cells. In such setting, cell type composition between samples were relatively homogeneous and usually it is unnecessary to perform data integration. When sample-driven variation is evident or in cases when dataset contains both 3’ and 5’ scRNA samples, to control against technical variations, Harmony^25^ were applied to datasets with (vars.to.regress = ‘chemistry’) option during data scaling. ‘Contaminant’ cells which mis-segregated into large pools were identified and put back into the unprocessed pool. Iterative processing of cells was done semi-automatically until all cells were processed. In the end, a total of 133953 cells (61724 epithelial, 5458 endothelial, 15203 fibroblast, 1918 myeloid, 23070 CD4, 12165 CD8, 783 NK and 13632 B cells) were collected for downstream study. Data were visualized with 2D tSNE or UMAP projection.

#### TCR integration

TCR sequencing data were processed with cellranger-tcr^59^ with standard parameters. After barcode matching and TCR de novo assembly, the assembled TCR sequence and clonotypes were extracted. Clones of cells with similar TCR in the same donor were identified as having exactly similar TCR exon usage combination as well as similar CD3r nucleotide sequence for all TCR (alpha and beta chain, if applicable) sequences expressed in the cell. Cells with ≥ 3 TCR sequences were removed. Clonal expansion is defined as a single T cell clone was observed >2 times in sequencing data from the donor. Local clonal expansion was defined as the clone is observed >2 times in one sample and <2 times in all other samples. Global clonal expansion was defined as the clone is observed >2 times in ≥2 samples. scRNA data were subsequently registered with this TCR information to annotate the clonal identity and clonal expansion status of each T lymphocyte.

#### Cell-cell signalling inference

Cell-cell signalling inference was done with CellChat^60^ with cells from normal bladder, bladder cancer tissue, adjacent bladder tissue and metastasis prostate tissue. Cell types with ≤ 20 cells were excluded from the analysis. Signalling probability was extracted from the object.

#### Co-regulated transcription module inference using NMF

Non-negative matrix factorization was performed in R using the iterative NMF (iNMF) internal function from Liger^61^ with data scaled by sample. After NMF, the weight matrix, variation matrix, and loading matrix were extracted from the object. Prediction of metagene expression in single cell was done with the top 100 contributors of NMF component using Seurat AddModuleScore function.

#### Transcription factor inference with cisTarget

RcisTarget^34^ was used to extract enriched TFBS in the co-regulated gene modules from NMF (top 100 genes from each component) with the human GRCh38 -1000bp∼+500bp database downloaded from cisTopic (hg38_refseq-r80_500bp_up_and_100bp_down_tss.mc9nr.feather). After extraction, high confidence co-regulator TF of regulons with NES>3.0 were extracted from the data. Specific TF regulating a NMF component was defined as a TF which was found in ≤ 2 components. TF binding affected by methylation was identified as the binding site is derived from the in vitro DNA-binding activity from Yin et al’s work^33^. Visualization of co-regulated transcription network is performed with visNetwork (https://cran.r-project.org/package=visNetwork). Matching of TF to cell-cell signalling event was done manually.

#### Developmental lineage analysis

Diffusion map of single cell RNA expression data was computed with destiny^62^. RNA velocity analysis of single cell 3’ RNA sequencing data was performed with scVelo^63^ using Reticulate^64^ in R (3.6.2) with Python 3. With RNA velocity defined the root cluster, slingshot was performed on diffusion maps to produce minimally spanning tree lineages.

#### Single cell ChrAcc data analysis

Single cell ATAC (chromatin accessibility, scATAC) raw reads were mapped with cellranger-atac^65^. The mapped fragment files were then processed by ArchR^30^. Quality control, doublet identification, LSI-based dimensionality reduction, cell clustering, gene expression inference, peakset identification (with MACS2), and marker peak finding were all performed in ArchR. Cells from all samples were pooled in the initial analysis, manually annotated with known markers, and separated into different subsets. The subset data were then subjected to scRNA integration in ArchR using Seurat CCA algorithm, with constrains for integration on large pools of cell type. scATAC signal of tumour epithelial cells (cancer cells) segregate into individuals because of apparent ‘chromatin accessibility’ change with CNV, which was impossible to be removed even using an integer cap to per-window reads. Hence, for epithelial cells, CNV analysis was done with the ‘tile’ fragments data with a circular binary segmentation algorithm. These regions were then intersected with CNV profiles of the same tumour from panel sequencing, and resulted ‘true’ copy number varied regions were used as a mask for a second round of LSI-based dimensionality reduction to perform copy-number-invariant clustering of cells. Peaks were called for single cell dataset using MACS2. Chromatin accessibility on given genomic regions was also calculated in ArchR for single cell chromatin accessibility difference analysis. Transcription factor footprinting (activity measurement) was done in ArchR with ChromVar. Co-accessible regions were calculated with ArchR with ‘addCoAccessibility’ (maxDist = 1000000) and filtered with correlation >0.1. To visualize scATAC coverage on given DMR regions, grouped single cell coverage files were written by ArchR and profile plots were made with deepTools ‘plotProfile’.

#### Epigenotype

MIBC-specific DMR and MIBC differential ATAC peaks were collected and merged. The merged ‘epigenetically change regions’ were then used as regions for dimentionality reduction with LSI in ArchR. Epigenotype clusters were then subjected to a further dimensionality reduction with diffusion map, and slingshot lineage inference. Epigenotype match to RNA cluster (phenotype) was counted in R. The Shannon entropy of epigenotype-phenotype match was calculated to infer the transcriptional plasticity of given epigenotype.

#### Cell cycle scoring

Single cell RNA cell cycle scoring was done with Seurat^26^ using the ‘cc.gene.2019’ dataset as source of G2/M and S-phase gene sets. For visualization, G2M.score and S.score of a single cell were added together to suggest the relative ‘actively cycling’ probability.

#### Differentiation potential assessment

Cellular differentiation potential were assessed by CytoTRACE^66^. Briefly, RNA expression matrix of single cells are extracted from Seurat object, with all transcripts regardless of their in-assay-variability. CytoTRACE analysis was performed on this matrix without downsampling. Per-cluster median CytoTRACE score was used as an index for the differentiation state of each cell cluster. For the comparison, we selected from the scRNA dataset all ‘normal’ urothelium cells, and cancer cells which projected to scATAC counterparts. Cancer scRNA cells were labelled by their projected scATAC counterparts.

#### SNV matching in single cell sequencing data

For mutation expression analysis, standard cellranger pipeline (https://support.10xgenomics.com/single-cell-gene-expression/software/downloads/latest) were used to produce a bam file from scRNA sequencing data. SNV were extracted from paired tumour mutation panel capture sequencing results and lifted-over to hg38 using easyLiftOver based on LiftOver. Only somatic mutations with >5% VAF were included in the analysis. Vartrix (https://github.com/10XGenomics/vartrix) was used to build a cell x mutation matrix (Supplementary Table 10).

#### Metagene activity in bulk RNA sequencing data

Metagene were defined as the top 100 genes that contribute to a NMF component. Raw read count matrix of bulk BLCA tumour tissue RNA sequencing were downloaded from UCSC XENA and IMVigor210 sources. Limma^67^ was used to normalize the RNAseq count matrix, and GSVA^68^ was used to score metagene activity. Patients/samples were classified as ‘high’/’low’ with respect to that metagene activity using a 2-mode Gaussian distribution fitting with mixtools^69^.

#### Survival analysis

Sample-centered clinical data were downloaded from UCSC XENA and IMVigor210 sources. Survival plots and analysis were done with R package survplot (http://www.cbs.dtu.dk/~eklund/survplot/).

## Data Availability Statement

All data, including raw data and clinical information, have been submitted to the Genome Sequence Archive for Human (http://bigd.big.ac.cn/gsa-human/) at the BIG Data Center, Beijing Institute of Genomics, Chinese Academy of Sciences, under the accession number HRA001225. The raw sequencing data and clinical information unique to an individual and require controlled access. The deposited and publicly available data are compliant with the regulations of the China Human Genetic Resources Management Office, Ministry of Science and Technology of China.

## Funding

This study was supported by grants from the Chinese Central Special Fund for Local Science and Technology Development of Hubei Province (2018ZYYD023), Research Fund of Zhongnan Hospital of Wuhan University (ZNJC201915, SWYBK, SWYBK01, SWYBK02), Science and Technology Department of Hubei Province Key Project (2018ACA159), and Improvement Project for Theranostic Ability on Difficulty Miscellaneous Disease (Tumor) from National Health Commission of China (ZLYNXM202006). The funders played no role in the study design, data collection and analysis, decision to publish, or preparation of the manuscript.

## Acknowledgments

The excellent technical assistance of Danni Shan, Shanshan Zhang, Wen Chen, Wan Xiang, Hongwei Peng and Zongning Zhou at ZNWH is gratefully acknowledged. We also would like to acknowledge the TCGA consortium and authors publicly sharing data used in our research. The authors disclose no writing assistance.

## Author Contributions

**Conceptualization**, Y.X., Y.Zhang, X.W.;

**Methodology**, Y.X., W.Jin, K.Q., K.W., H.Z., Y.W., F.C., Y.Zhang;

**Computation and Formal analysis**, W.Jin, K.W., L.S., F.C., Y.Zhang;

**Investigation**, Y.X., W.Jin, K.Q., K.W., G.W., L.J., Y.Zhao, L.C., X.Chen, N.L., L.S., Sheng Li, H.S., M.P., Y.Zhang;

**Resources**, K.Q., G.W., R.C., L.J., Y.Zhao, H.Z., T.L., Z.X., T.W., J.L., F.Y., D.L., H.C., Z.Y., Sheng Li, H.S., Z.G., Y.G., N.L., S.J.L., W.D., W.Z., D.C., Y.T., Y.L., C.J., X.H., H.M., X.W.;

**Writing Original Draft, Review, and Editing**, Y.X., W.Jin, K.Q., G.W., W.Jiang, R.C., L.J., Y.Zhao, X.Cao, M.P., Y.Zhang, X.W.;

**Writing Critical Review**, Commentary or Revision, Y.X., K.Q., G.W., W.Jiang, L.J., Y.Zhang, X.W.;

**Writing Data Presentation**, Y.X., W.Jin, K.Q., G.W., W.Jiang, L.J., Y.Zhang;

**Supervision**, Z.X., Y.Zhang, X.W.;

**Project Administration**, Y.X., K.Q., Z.X., Shenjuan Li, Y.Zhang, X.W.;

**Funding acquisition**, Y.X., Y.Zhang, X.W.

## Competing interests

The authors declare that they have no competing interests.

## Notes

http://bigd.big.ac.cn/gsa-human/

### Competing Interest Statement

The authors have declared no competing interest.

