## Supplementary information for "Emergence of an adaptive epigenetic cell state in human bladder urothelial carcinoma evolution"

### Supplementary Notes

#### Tumour microenvironment of BLCA

Tumour-infiltrated cytotoxic T cells include putative tumour-reactive CD8 ($CD39^{+}$(*$ENTPD1^{+}$*)/$CD103^{+}$(*$ITGAE^{+}$*)/*$PDCD1^{+}$*), bystander CD8 ($CD39_{low}$/$CD103_{low}$/*$GZMK^{+}$/$TOX^{+}$*), and exhausted CD8 (*$EOMES_{hi}$/$TBX21_{hi}$/$IFNG_{low}$*) (Fig. 1H and Supplementary Fig. 7E). The prevalence of activated CD8 (IFNG+) in tumour tissue was increased as clinical stage progress (Fig. 1N and Supplementary Fig. 7J). Tumour-infiltrated cytotoxic T cells, either bystander or tumour-reactive, were identified in the majority of T3-T4 BLCA; while exhausted CD8 T cells were relatively rare in our samples (Fig. 1N and Supplementary Fig. 7J). A tumour-reactive CXCR6-positive cytotoxic T-cell population ($trT_{eff}$ CXCR6) is distinct from other $trT_{eff}$ cells by overexpressing *CXCL13/TGFBR3/CEMIP2*(migration associated gene)/*PBX4* and cross-tissue clonal distribution (Supplementary Fig. 7). Prevalence of $trT_{eff}$ CXCR6 was significantly associated with longer survival in immunocheckpoint inhibitor (ICI)-treated patients (Supplementary Fig. 7), suggesting a pivotal role for it in anti-PD1 dependent anticancer responses.

The tumour-infiltrated CD4 T helper cells consisted of activated Th1, Th17, Treg and bystander (bs Th2/17) populations (Fig. 1I and Supplementary Fig. 6B). Putative tumour-reactive Th1 (trTh1, *PDCD1+/TBX21+/LAG3+/IFNG+*/CD39hi/CD103hi) and tumour-reactive Treg (trTreg, *FOXP3+/CD274+/LAG3+/TBX21+*/CD39hi/CD103hi, overexpressing *CD274*/*TNFRSF4* /*TNFRSF18*/*TNFAIP3*) could be identified in the tumour-infiltrated population (Supplementary Fig. 6). Tumour reactiveness of these cells is not only marked by CD103/CD39 expression but also local TCR clonal expansion. Patients with advanced stage disease were associated with increased prevalence of infiltrated tumour-reactive Treg and bystander Th2/17 cells (*PDCD1+/CD200+/NRP1+*/CD39-/CD103-) (Fig. 1N and Supplementary Fig. 6). In concordance to the previous reports[1], all tumour-reactive $CD4^{+}$ T populations were locally expanded, as cells of the same TCR clone are only observed in tumour tissue but not in other tissues from the same donor. Trajectory analysis of $CD4^{+}$ T lineage suggested that trTreg and eTreg bifurcated early (Supplementary Fig. 6). In ICI-treated patients, eTreg level was negatively associated with survival time, whilst the presence of both trTreg and $trT_{eff}$ CXCR6 predicted a longer survival (Supplementary Fig. 6M).

The prevalence of NK cells (*NKG7*$^{+}$/*XCL2*$^{+}$/*CD3D*$^{-}$) is relatively few in our samples (Fig. 1C) and could be classified into immature (*IL7R/TGFB1*), proliferating (*TOP2A/ZEB2/TBX21*), mature cytotoxic (*GZMH/GZMM*), and mature post-activation NK (*EOMES/HAVCR2/INFG*) (Fig. 1J and Supplementary Fig. 9). B-lymphocytes, characterized by *CD79A/MS4A1*, are only found in T1-T4 stages, but not Ta-stage tumour or healthy bladder (Fig. 1K, N and Supplementary Fig. 10). Myeloid cells, characterized by *LYZ*, were classified into monocytes, classical dendritic cells (cDC: *CLEC9A/CD1C*) and macrophage (*CSF1R/CD163*) (Fig. 1L and Supplementary Fig. 11). For NK, B and myeloid cells, we did not observe an obvious association with BLCA clinical stage.

Consistent with previous report[2], different subtypes of fibroblasts were identified between tumour and cancer-adjacent tissues (Fig. 1M). DCN$^{+}$ population is only found in normal or adjacent bladder tissue (Supplementary Fig. 12). Cancer-associated fibroblast, both *RGS5*$^{+}$/*JAG1*$^{+}$ and *PDGFRA*$^{+}$/*MMP2*$^{+}$ populations, might contribute to tumour induction and tissue maintenance[2].

**Luminal and basal subtypes in human BLCA**

Clinical observations show patients with superficial papillary BLCA are more likely to recur, but only a fraction of them progress to highly invasive BLCA. Meanwhile, most of the high-grade invasive BLCA patients do not have a history of superficial papillary BLCA[3]. Therefore, two distinct tracks, termed papillary/luminal and nonpapillary/basal tracks, are postulated for BLCA progression[4,5]. However, the initiating events of these two tracks are not revealed and which cell types they may originate from is not identified[6]. Muscle-invasive cancer cells are likely originated from the basal urothelium in an animal model of MIBC induced by N-butyl-N-(4-hydroxybutyl) nitrosamine (BBN)[7]. In another study, the progenitors in NMIBC tumours were speculated to originate from non-basal cells[8]. In human, immunohistochemistry (IHC) analysis of clinical BLCA sections revealed expression of CK5/6 in MIBC and CK20 in NMIBC, similar to the finding in mouse. However, no direct evidence was made from molecular level to confirm that basal/luminal types of human BLCA originates from distinct celltypes. Our results show that in human, MIBC originate from basal cells and ancestors of NMIBC are superficial intermediate/umbrella cells. More specifically, our analysis of single cell gene expression and tissue methylation data suggested that the cancer cells originated from different basal cell lineage processing different degree of malignancy, whilst the ones originated directly from BsP being most malignant. As basal/luminal types of bladder cancer resembled the classes in breast cancer, whether different classes of breast cancer originate from different progenitors is an open question.

**Function of TM4SF1**

TM4SF1, a member of the tetraspanin family, has been identified as a tumour-specific antigen, promoting proliferation, invasion, epithelial-mesenchymal transition (EMT) and chemo-resistance[9-11]. As a surface antigen and associated with pathologic angiogenesis, TM4SF1 is considered as a potential therapeutic target for specific monoclonal antibodies or drug-conjugated antibodies[12,13]. In addition, Bae and colleagues reported that TM4SF1 could discriminate mesenchymal stem cells (MSCs) simultaneously from blood cells and fibroblasts[14]. Our previous study found that TM4SF1 is upregulated in human MIBC tissues, and associated with T stage, lymph node metastasis status and survival rate of MIBC patients[15]. Moreover, TM4SF1 could regulate apoptosis, cell cycle and ROS metabolism, through PTEN/PI3K/AKT pathway and WNT/β-catenin pathway, the main driver of the aggressive invasive phenotype of BLCA[5,11]. Here, we identified that TM4SF1 also marks human bladder urothelium progenitor cell (Fig. 2) as well as the hyperplastic cancer-stem-like TPCS. These results echo the recent human lung study[16] and suggest that TM4SF1 might serve as a general stem cell marker in many tissues with layered epithelium structures. The exact function of TM4SF1 in these cells remains to be an open question.

**The influence of ITH on immunogenicity of BLCA**

Besides cancer cells, the tumour microenvironment (TME) consists of various immune cells, cytokines, stromal cells, and related factors they secreted. The TME composition is associated with the prognosis and therapeutic effect of cancer patients[17]. BLCA is known to be immunogenic and can respond to BCG and immune checkpoint inhibitors[18]. However, intravesical BCG treatment is not working in 25% - 45% of BLCA patients, and nearly 40% of patients receiving BCG treatment will eventually relapse[19]. In addition, the overall response rate of PD1/PD-L1 inhibitors is not satisfactory for the treatment of advanced BLCA[18]. Therefore, understanding the immunological characteristics of the tumour microenvironment is emerging required for developing beneficial treatment strategies for patients with BLCA. Studies found that the ratio of $CD8^+$ T cells over myeloid-derived suppressor cells is a predictive biomarker of recurrence-free survival in NMIBC patients[20]. Intratumoral $TIGIT^+CD8^+$ T cells and the ratio of $CD8^+$ T cells to $T_{reg}$ cells in the pre-treatment BLCA tissues are associated with response to neoadjuvant chemotherapy[21,22]. Metastatic BLCA patients with intense TILs receiving platinum-based chemotherapy indicate prolonged overall survival[23], and those with higher TMB and enrichment of $CD8^+$ T effector cell have a higher objective response rate to atezolizumab[24]. In addition, tumour cells and $Lamp3^+$ DC subsets can cooperate with each other to reduce immunogenicity and form an immunosuppressive microenvironment by down-regulating MHC-II molecules[2]. These results revealed a complex landscape of immune-tumour interaction in BLCA. We found that with clinical stage progress of BLCA, the activated tumour associated cytotoxic $CD8^+$ T cells, containing numerous $CD39_{low}/CD8^+$ and $CD39_{low}/CD8^+$ bystander T cells, and helper $CD4^+$ T cells are increased in tumour tissues. Even in late stage tumour, the clonal expansion of T cells is still active, and naïve T cells are continuously recruited and become tumour-reactive T cells in BLCA. Anti-PD1 treatment is found to be most effective in tumours enriched with both trTcyt and trTreg cells, suggesting that trTreg-initiated PD-L1/PD-1 interaction is the prevalent immunosuppressive mechanism in these tumours. TPCS is found to be shifting between mutation-expressing and mutation-free RNA expression states, suggesting a 'digital'-like dynamic transcriptional regulation similar to those observed in synthetic and natural repressilator gene networks. Furthermore, TPCS prevalence is correlated with immune infiltration and T stage progression. Together, our results suggested not only how tumour-reactive cytotoxic T cell population is being regulated by other cells in the TME, but also how cancer cells change after immunoediting to escape from tumour-reactive cytotoxic T cell attack.

**The influence of ITH on chemotherapy resistance of BLCA**

Currently, cisplatin-based combination chemotherapy is still the first-line treatment for metastatic BLCA[25], but faces the challenge of therapy resistance. Zhang and colleagues described the existence of CSCs in cisplatin-resistant bladder cancer cells and suggested that Nanog and Bmi1 may play an important role in malignancy[26]. Another report suggested that the COX2/PGE2 pathway and the YAP1 pathway co-recruit SOX2 expand the urinary CSCs population and maintain chemotherapy resistance[27]. More recently, Hsu and colleagues demonstrated that the EMT upregulates the expression of PD-L1 via β-catenin/STT3 signalling axis in CSC, thereby contributing to CSC immune evasion[28]. Here, we report that the prevalence of TPCS strongly increases in the patients receiving gemcitabine intravesical in our cohort. In contrast to all other cancer cell types which entered a dormant-like state post chemotherapy, cell proliferation ability of TPCS was not as sensitive to chemotherapy. Analyzing the RNA expression profiles from BLCA patients revealed the association between higher TPCS markers expression and worse prognosis after chemotherapy. These results suggest that the cancer stem-like cell TPCS might function as the key mediator of chemotherapy resistance.

#### Supplementary Figures

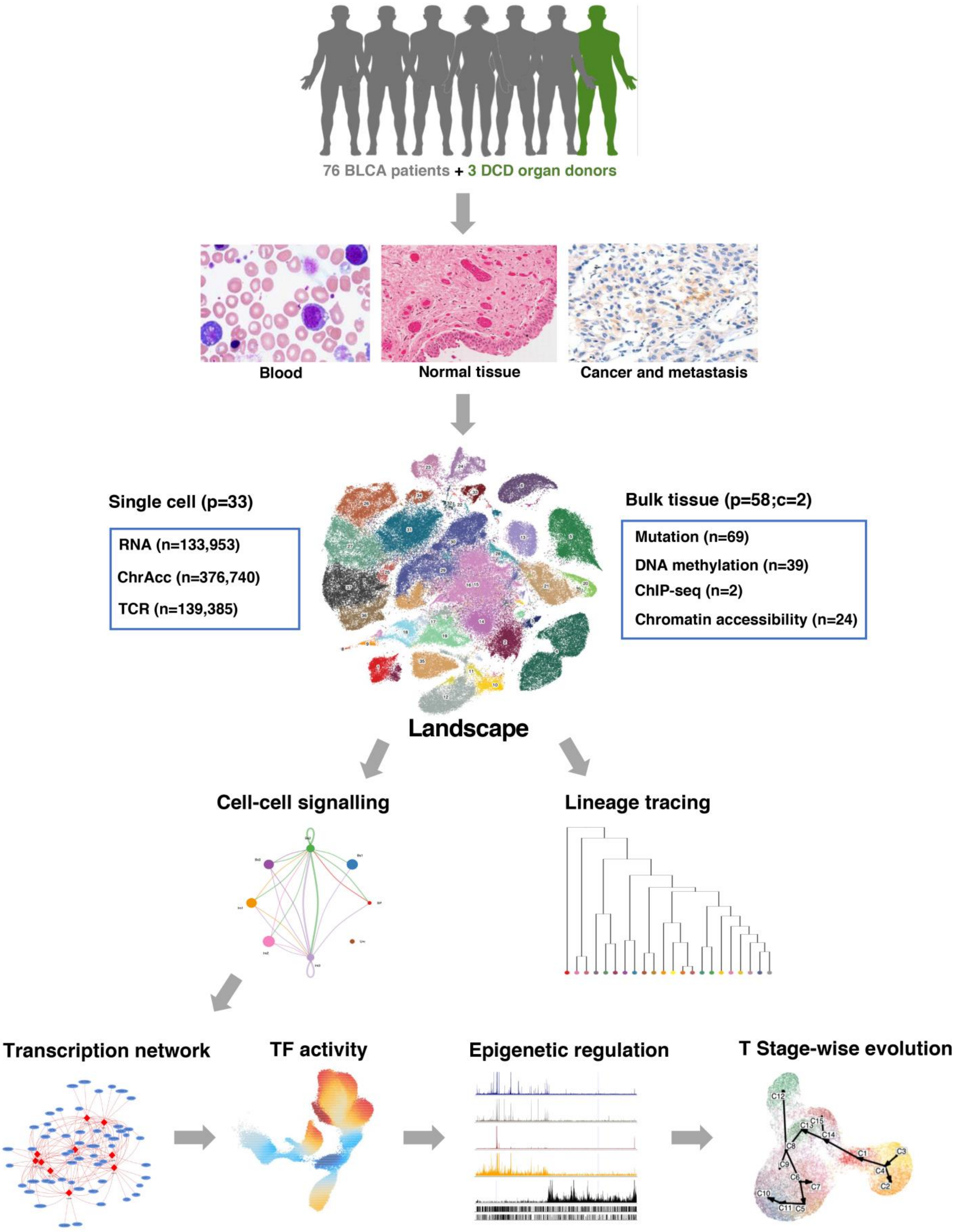

**Supplementary Figure 1. Schematic representation of experimental strategy for the study.**

To understand the tumour microenvironment evolution during bladder cancer progression across clinical stages and therapy, we performed a multi-center human study. A total of 65 BLCA patients and 3 Donation after Cardiac Death (DCD) controls were collected from 4 centers. Genomic mutation, DNA methylation, histone modification, transcription factor binding, chromatin accessibility, RNA expression and TCR rearrangement were profiled for bulk tissue from 2 cell lines and 58 persons (55 BLCA patients and 3 DCD controls; p: person; c: cell line) or single cell samples from a total of 33 persons (30 BLCA patients and 3 DCD controls). Additionally, 11 BLCA scRNA sequencing data were collected from public repository. In total, our collection included 122 tissues from 79 donors.  These tissue samples result in 59 3' scRNA libraries from 33 donors, 39 5' scRNA libraries from 20 donors, 32 scTCR libraries from 17 donors, 22 scATAC libraries from 22 donors, 24 bulk tissue ATAC libraries from 24 donors, 39 bulk tissue DNA methylation libraries from 31 donors, and 69 mutation profiling libraries from 51 donors. Meanwhile, bioinformatic analysis including cell-cell signalling analysis, transcription network construction, transcription factor activity measurement and epigenetic regulation analysis were performed in our study.

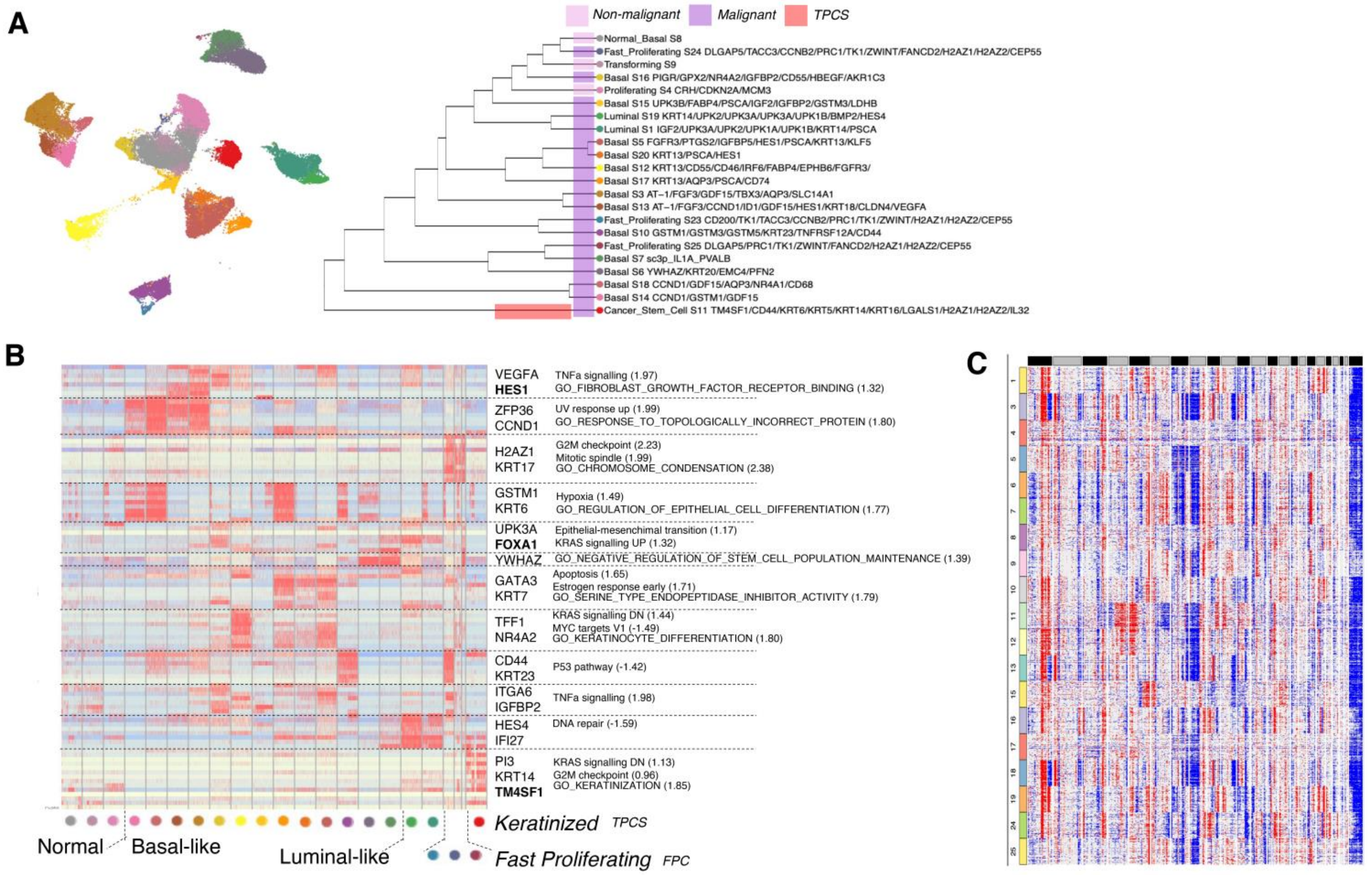


**Supplementary Figure 2. Epithelial scRNA analysis.**

**(A)** Left: UMAP-projection of single cell RNA sequencing with epithelial cells from bladder tissues. Right: Unsupervised hierarchical clustering with RNA expression. **(B)** Marker gene expression and enriched signalling pathway for epithelial clusters in (A). **(C)** Copy number analysis among epithelial clusters.

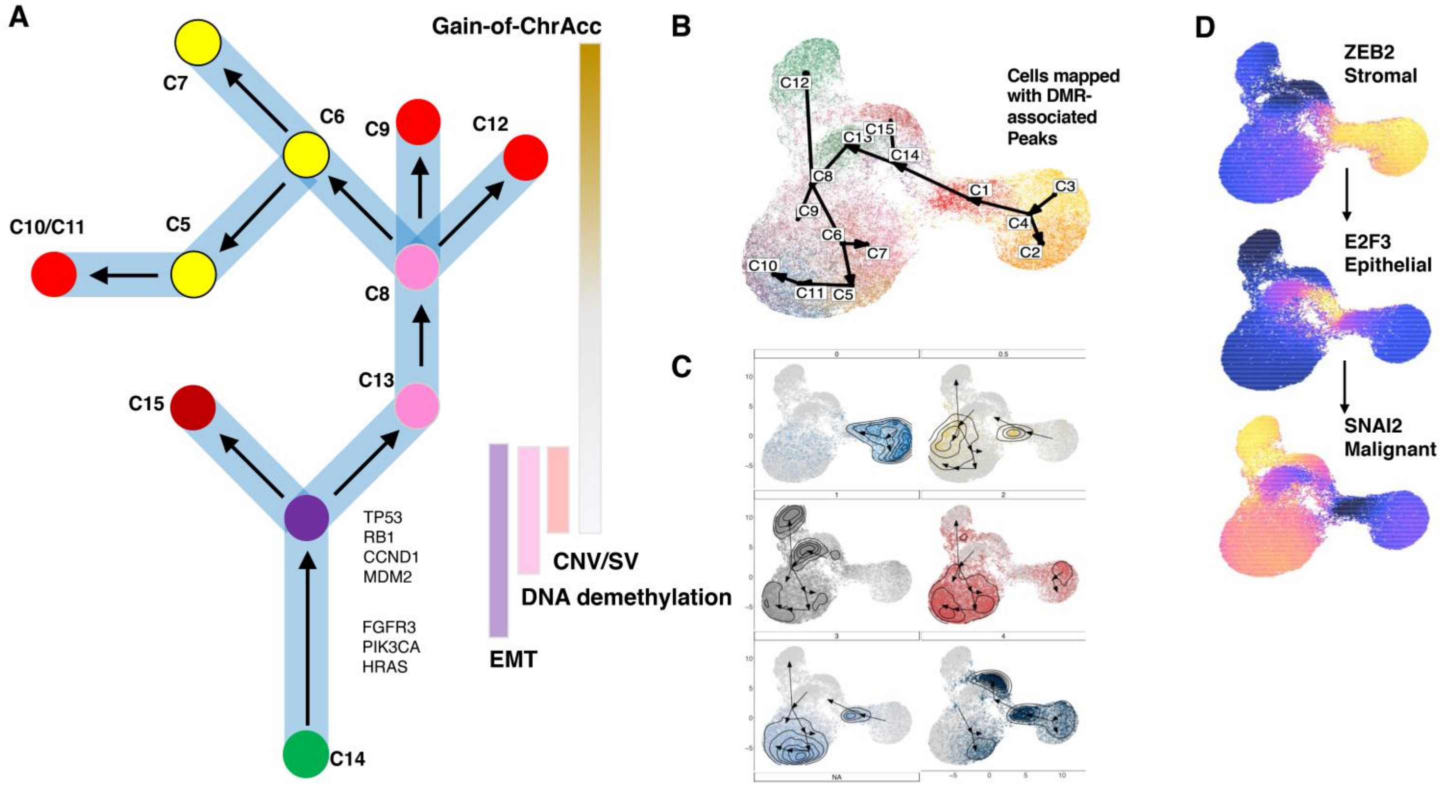


**Supplementary Figure 3. Epigenotype lineage development in scATAC cells.**

**(A)** Phylogeny diagram of bladder cancer evolution from the normal (C14) epithelial cells towards cancer. Molecular changes are shown on the right of the diagram. TPCS clusters are labelled in yellow (C5/6/7). Basal-like cancer cell clusters are labelled in red. Luminal-like cancer cell cluster (C15) is in dark red. Ta-associated, early cancer cell clusters are in pink. **(B)** UMAP projection of single scATAC cells overlaid with slingshot trajectory. **(C)** Distribution of cells in different stages from 0 (normal DCD) to 0.5 (Ta), and T1-T4 (1-4). Evolution trajectories of these cells are shown to reveal their origin (the begin of arrow) and fate (end of arrow). We noticed a significant switch between C10/C11 towards C5/6/7 branch during clinical stage progression. **(D)** Expression of marker gene ZEB2, E2F3 and SNAI2 in scATAC cells, showing their respective identity.

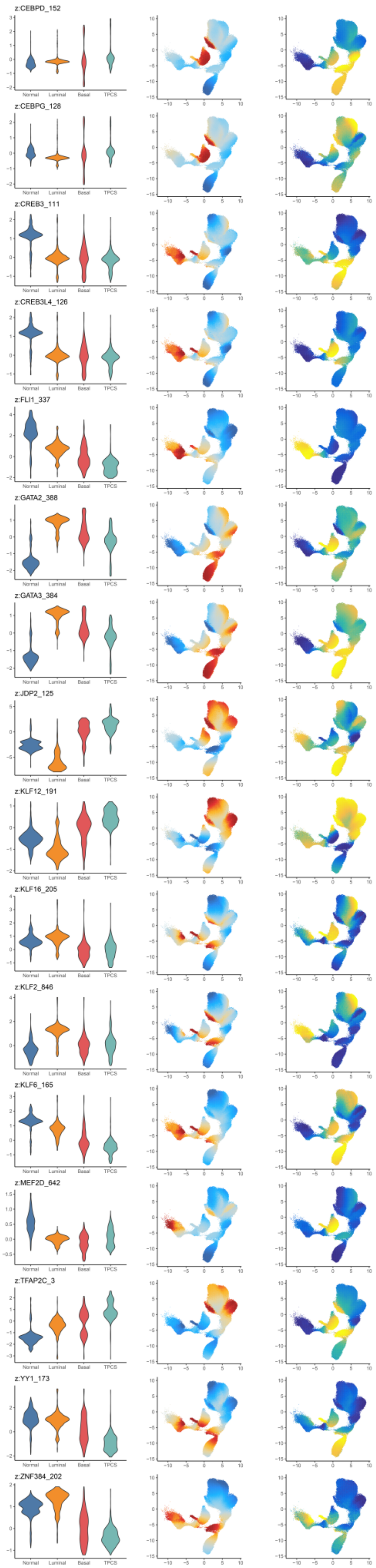
z:CEBPD_152
z:CEBPG_128
z:CREB3_111
z:CREB3L4_126
z:FLI1_337
z:GATA2_388
z:GATA3_384
z:JDP2_125
z:KLF12_191
z:KLF16_205
z:KLF2_846
z:KLF6_165
z:MEF2D_642
z:TFAP2C_3
z:YY1_173
z:ZNF384_202
Normal
Luminal
Basal
TPCS

**Supplementary Figure 4. Methylation-sensitive, significantly deviated TF activities in TPCS.**

From left to right: **(Left)**: Z-normalized genome-wide transcription factor binding activity on given TFBS motif measured in scATAC, grouped by classes of epithelial cells (normal, luminal (luminal-like cancer), basal (basal-like cancer), and TPCS (TM4SF1 positive cancer subpopulation)). **(Middle)**: Z-normalized genome-wide transcription factor binding activity on given TFBS motif measured in scATAC, projected to the UMAP manifold and smoothened by MAGIC in ArchR. **(Right)**: deduced gene expression level of given transcription factor in scATAC, projected to the UMAP manifold and smoothened by MAGIC in ArchR. Data shown in this figure are transcription factors that: 1. Showed significant DNA-binding activity deviation between TPCS and other types of epithelial cells; 2. Showed differential expression between TPCS and other types of epithelial cells; 3. Whose binding site is enriched in promoters of TPCS-specific NMF metagene. 4. The binding site of (3) is reported to be methylation-sensitive33.

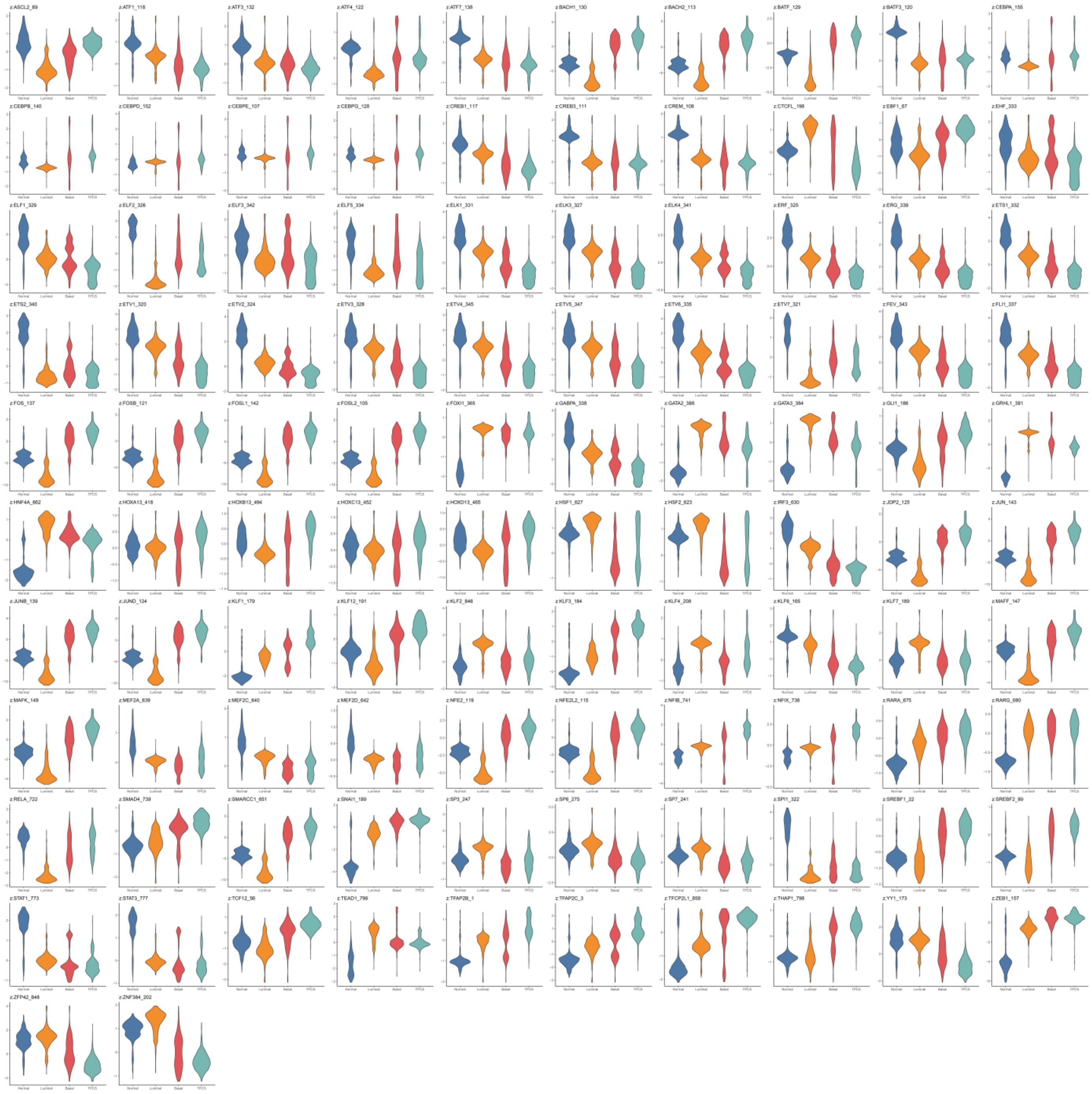

**Supplementary Figure 5. Methylation-insensitive, significantly deviated TF activities in TPCS.**

Each panel: Z-normalized genome-wide transcription factor binding activity on given TFBS motif measured in scATAC, grouped by classes of epithelial cells (normal, luminal (luminal-like cancer), basal (basal-like cancer), and TPCS (TM4SF1 positive cancer subpopulation)). Data shown in this figure are transcription factors that: 1. Showed significant DNA-binding activity deviation between TPCS and other types of epithelial cells; 2. Whose binding site is enriched in promoters of TPCS-specific NMF metagene. 3. The binding site of (2) is unknown to be methylation-sensitive33.

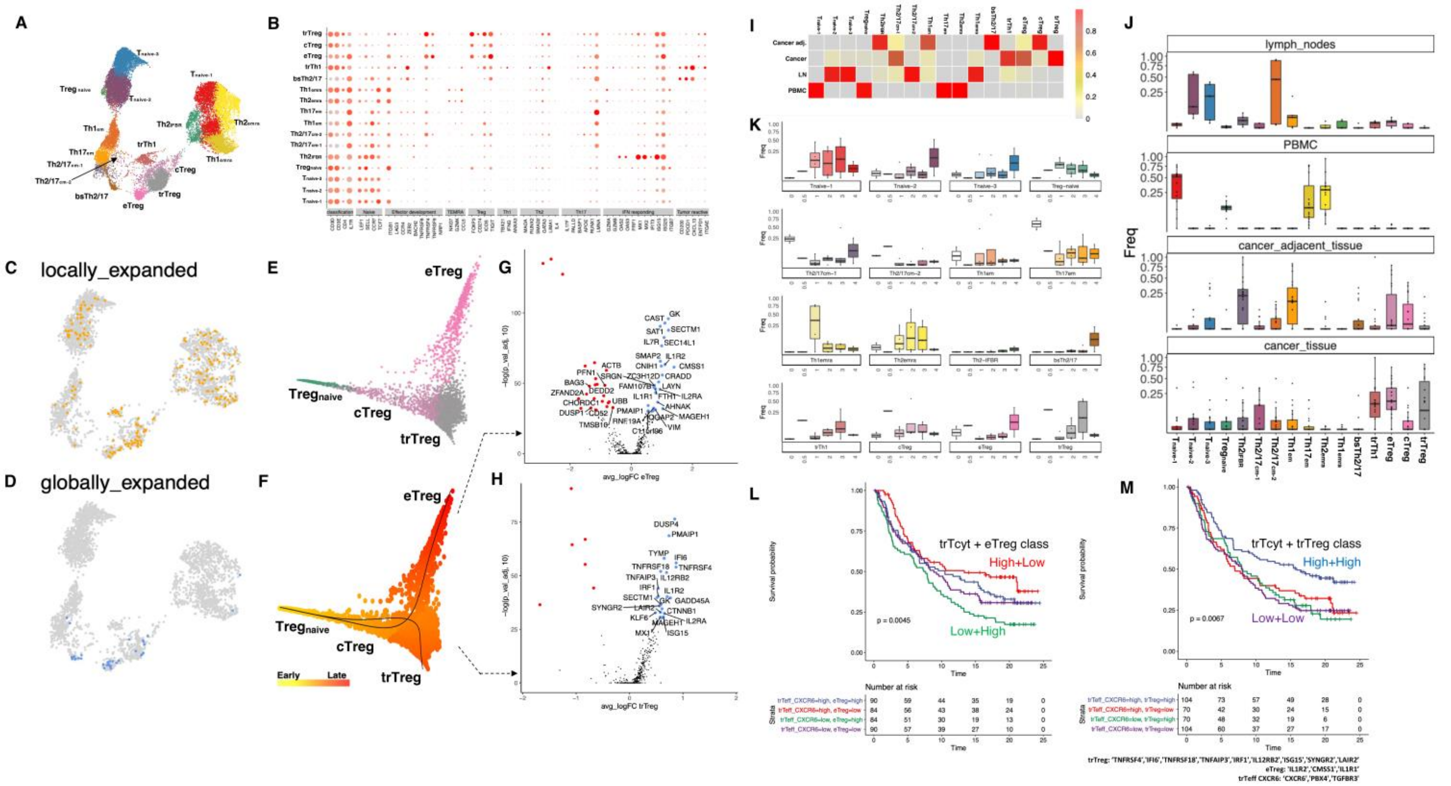


**Supplementary Figure 6. CD4$^{+}$ scRNA analysis.**

**(A)** UMAP-projection of single cell RNA sequencing with CD4 T cells from tissue and PBMC. **(B)** Marker gene expression for CD4 T cell clusters in (A). **(C)** UMAP-projection of locally expanded CD4 T cells. **(D)** UMAP-projection of globally expanded CD4 T cells. **(E)** Diffusion map projection for $T_{reg}$ in (A). **(F)** Developmental trajectory and pseudotime imputation of $T_{reg}$. **(G)** Differentially expressed genes between c$T_{reg}$ and e$T_{reg}$. **(H)** Differentially expressed genes between c$T_{reg}$ and tr$T_{reg}$. **(I)** Heatmap of tissue distribution of CD4 T cell types in (A). **(J)** Prevalence of CD4 T cell types in different tissues. **(K)** Prevalence of CD4 T cell types in (A) from healthy donor (0) to T4 stage muscle invasive bladder cancer donor (4). "0.5" denotes Ta. **(L)** Survival probability of patients with different tr$T_{eff\ CXCR6}$ and e$T_{reg}$ metagene activity in IMvigor210 cohort. **(M)** Survival probability of patients with different tr$T_{eff\ CXCR6}$ and tr$T_{reg}$ metagene activity in IMvigor210 cohort.

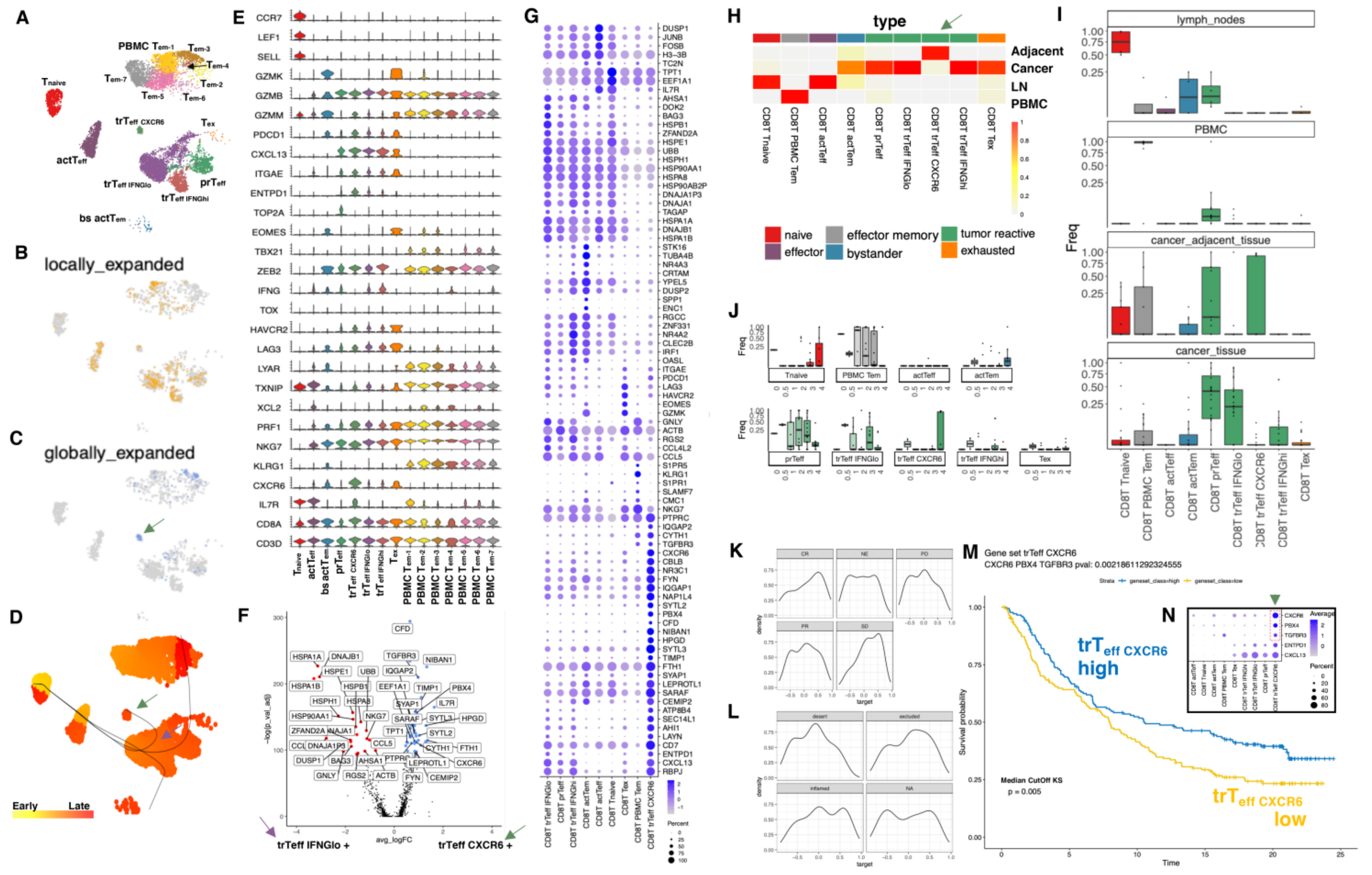


**Supplementary Figure 7. CD8$^{+}$ scRNA analysis.**

**(A)** UMAP-projection of single cell RNA sequencing with CD8 T cells from tissue and PBMC. **(B)** UMAP-projection of locally expanded CD8 T cells. **(C)** UMAP-projection of globally expanded CD8 T cells. **(D)** Development trajectory and pseudotime of CD8 T cells. **(E)** Marker gene expression for CD8 T cell clusters in (A). **(F)** Differentially expressed (DE) genes between trT$_{\text{eff CXCR6}}$ and trT$_{\text{eff IFNGlow}}$. Blue represents high expression in trT$_{\text{eff CXCR6}}$ and red indicates elevated expression in trT$_{\text{eff IFNGlow}}$. **(G)** Expression of DE genes between trT$_{\text{eff CXCR6}}$ and trT$_{\text{eff IFNGlow}}$ in total CD8 T cell clusters in (A). **(H)** Heatmap of tissue distribution of CD8 T cell type in (A)**. (I)** Prevalence of CD8 T cell types in (A) in different tissues. **(J)** Prevalence of CD8 T cell types in (A) from healthy donor (0) to T4 stage muscle invasive bladder cancer donor (4). "0.5" denotes Ta. **(K)** Distribution of trT$_{\text{eff CXCR6}}$ metagene activity in patients with different responses (CR, PR, SD, PD, and NE) to Atezolizumab in IMvigor210 cohort. **(L)** Distribution of trT$_{\text{eff CXCR6}}$ metagene activity in tumours with different immune infiltration type (inflamed, excluded, and immune-desert) in IMvigor210 cohort. **(M)** Survival probability of patients with high and low $trT_{eff\ CXCR6}$ metagene activity in IMvigor210 cohort. **(N)** Expression of $trT_{eff\ CXCR6}$ metagene in CD8 T cell clusters in (A).

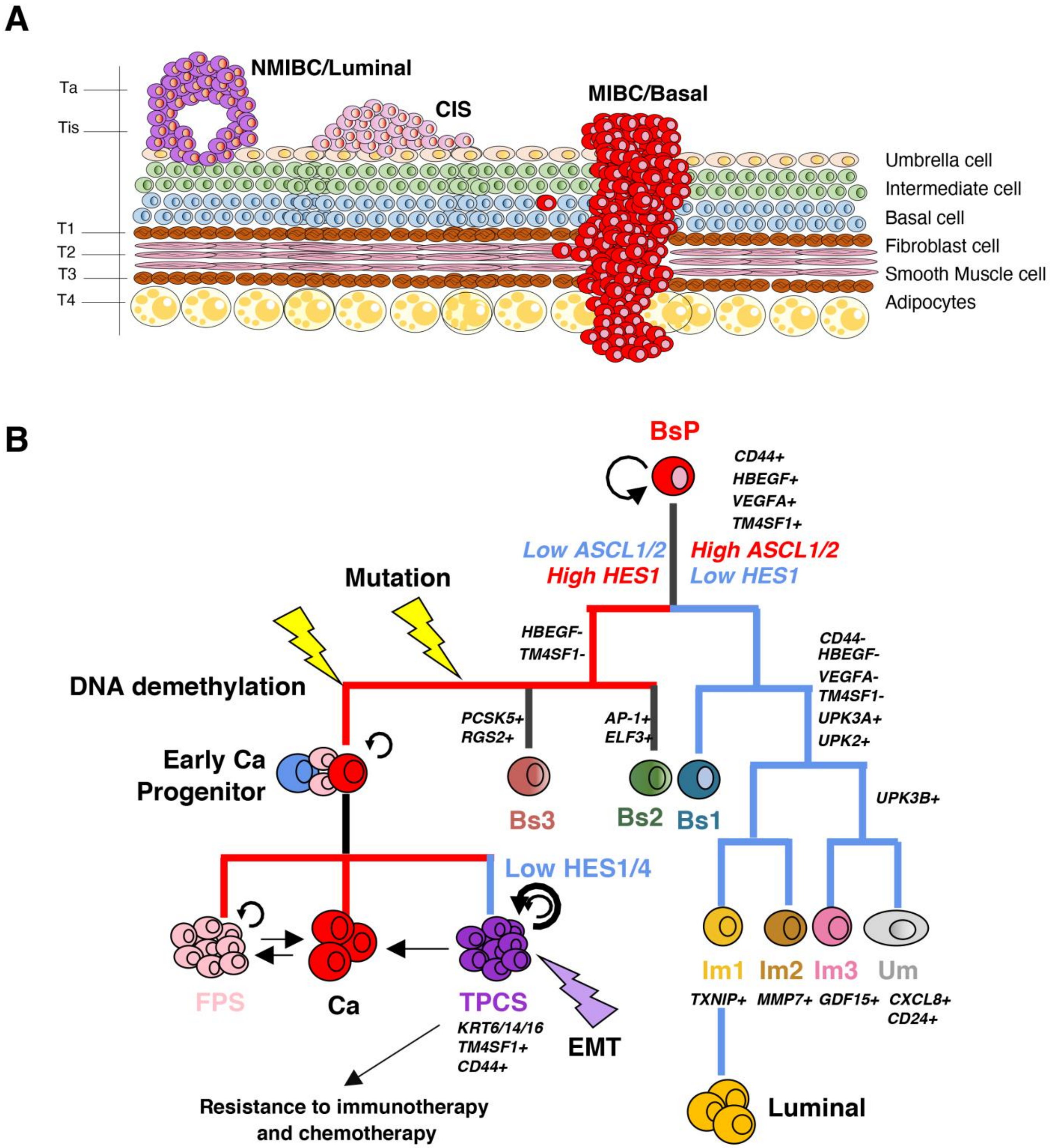


**Supplementary Figure 8**. **Summary of the study.**

**(A)** Schematics summarizing bladder cell types, the clinical stages and subtypes of BLCA. **(B)** The diagram of the evolution of BLCA in our study, revealing a step-wise pathway for the emergence of a transcriptionally plastic, clinically important TM4SF1-positive cancer cell subpopulation (TPCS).

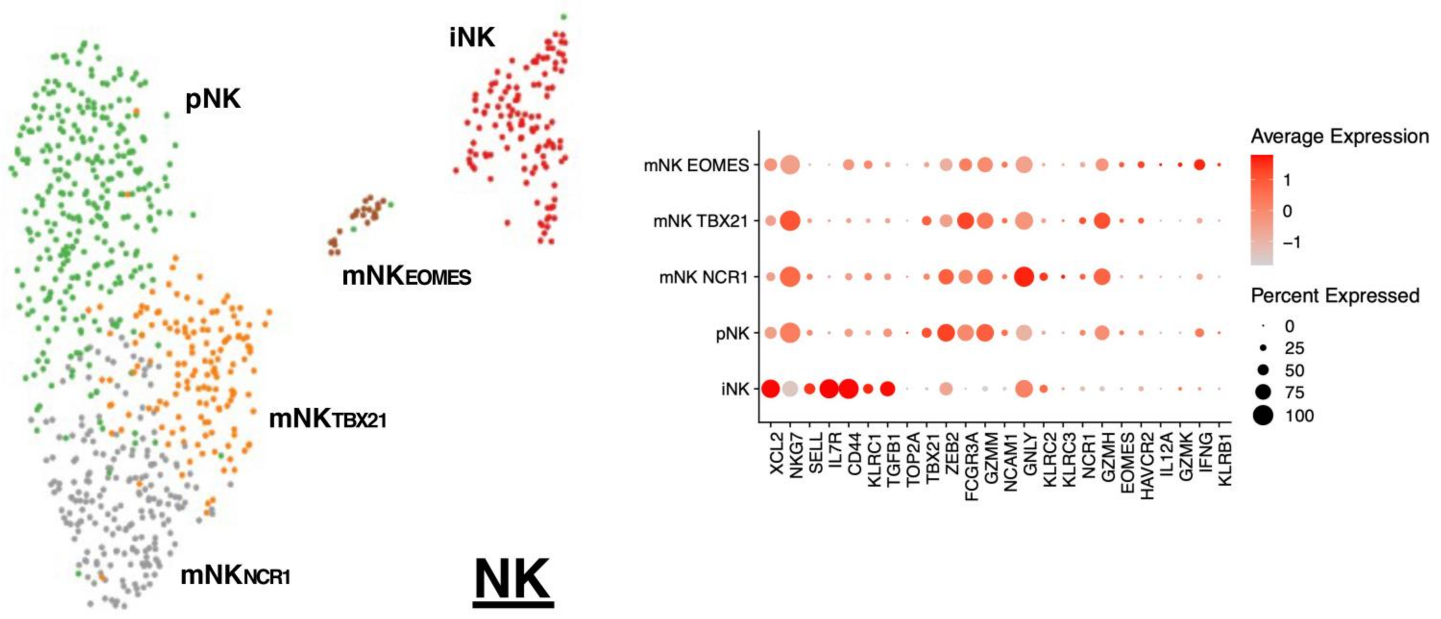


**Supplementary Figure 9. NK scRNA analysis.**

**(Left):** UMAP-projection of single cell RNA sequencing with NK cells from tissue and PBMC. iNK: immature NK; pNK: proliferating NK; mNK: mature NK. **(Right):** Marker gene expression in different NK clusters.

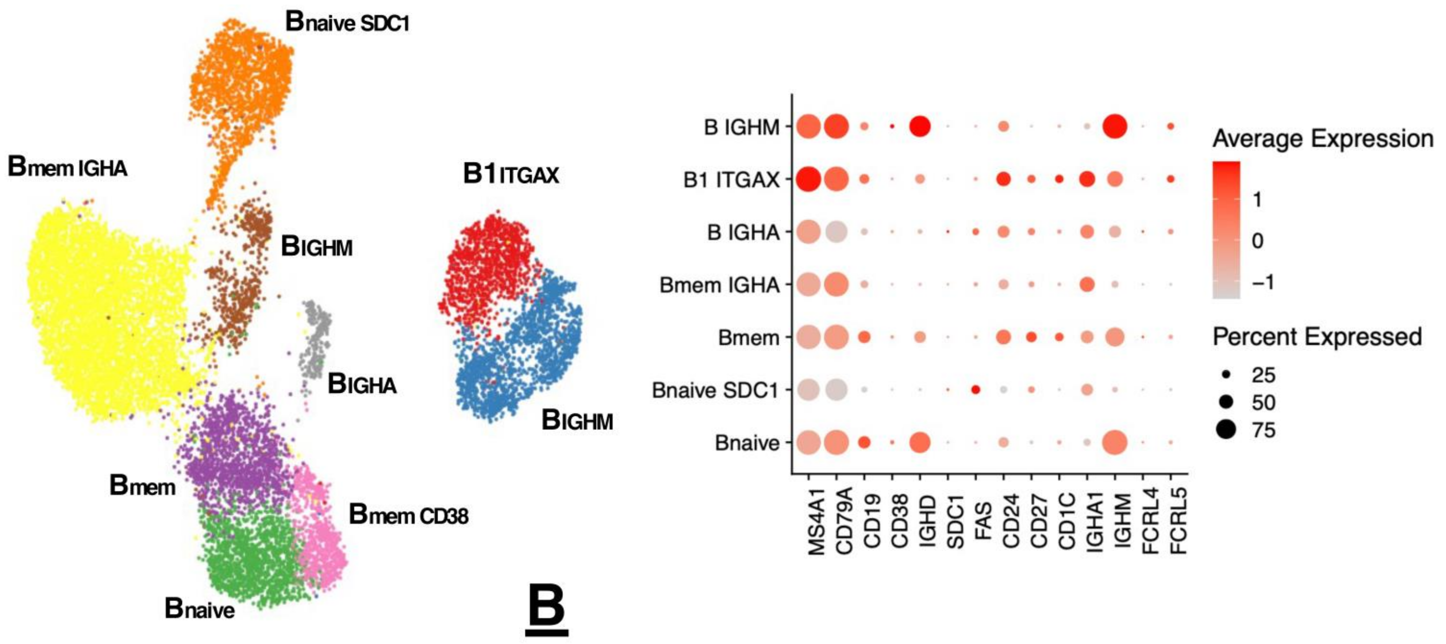


**Supplementary Figure 10. B lymphocyte scRNA analysis.**

**(Left):** UMAP-projection of single cell RNA sequencing with B cells from tissue, lymph nodes, and PBMC. $B_{\text{naïve}}$ SDC1: unswitched B without IGHM expression; $B_{\text{naïve}}$: IGHM-expressing, unswitched B; $B_{\text{mem}}$: memory B; $B_{\text{IGHM}}$: IGHM/IGHD expressing B; $B_{\text{IGHA}}$: IGHA expressing B. **(Right):** Marker gene expression in different B cell clusters.

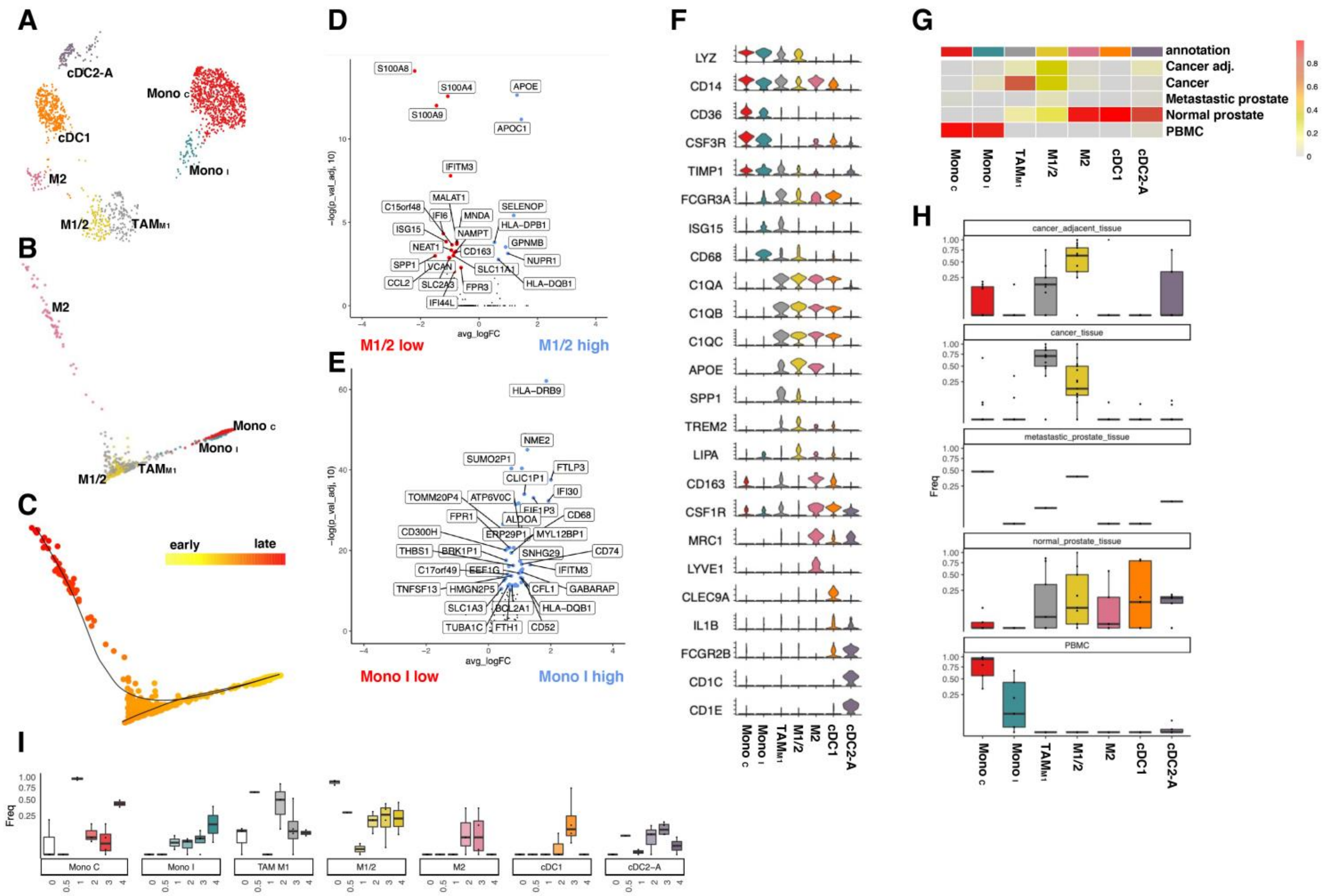


**Supplementary Figure 11. Myeloid scRNA analysis.**

**(A)** UMAP-projection of single cell RNA sequencing with myeloid (macrophage, monocyte, and dendritic) cells from tissue and PBMC. TAM: tumour-associated macrophage; Mono c: classical monocyte; Mono i: intermediate monocyte; cDC: classical dendritic cell. **(B)** Diffusion map of monocyte-to-macrophage lineage; **(C)** Slingshot trajectory built upon monocyte-to-macrophage lineage diffusion map, coloured by early (yellow) to late (red) pseudotime of differentiation. **(D)** Differentially expressed genes (x: logFC, y: -logPadj) between M1/2 and TAM$_{M1}$. **(E)** Differentially expressed genes (x: logFC, y: -logPadj) between Mono I and Mono C. **(F)** Marker gene expression in myeloid cells. **(G)** Heatmap of tissue distribution of myeloid cell type in (A)**.** (**H)** Prevalence of myeloid cell types in (A) in different tissues. **(I)** Prevalence of myeloid cell types in (A) from healthy donor (0) to T4 stage muscle invasive bladder cancer donor (4). “0.5” denotes Ta.

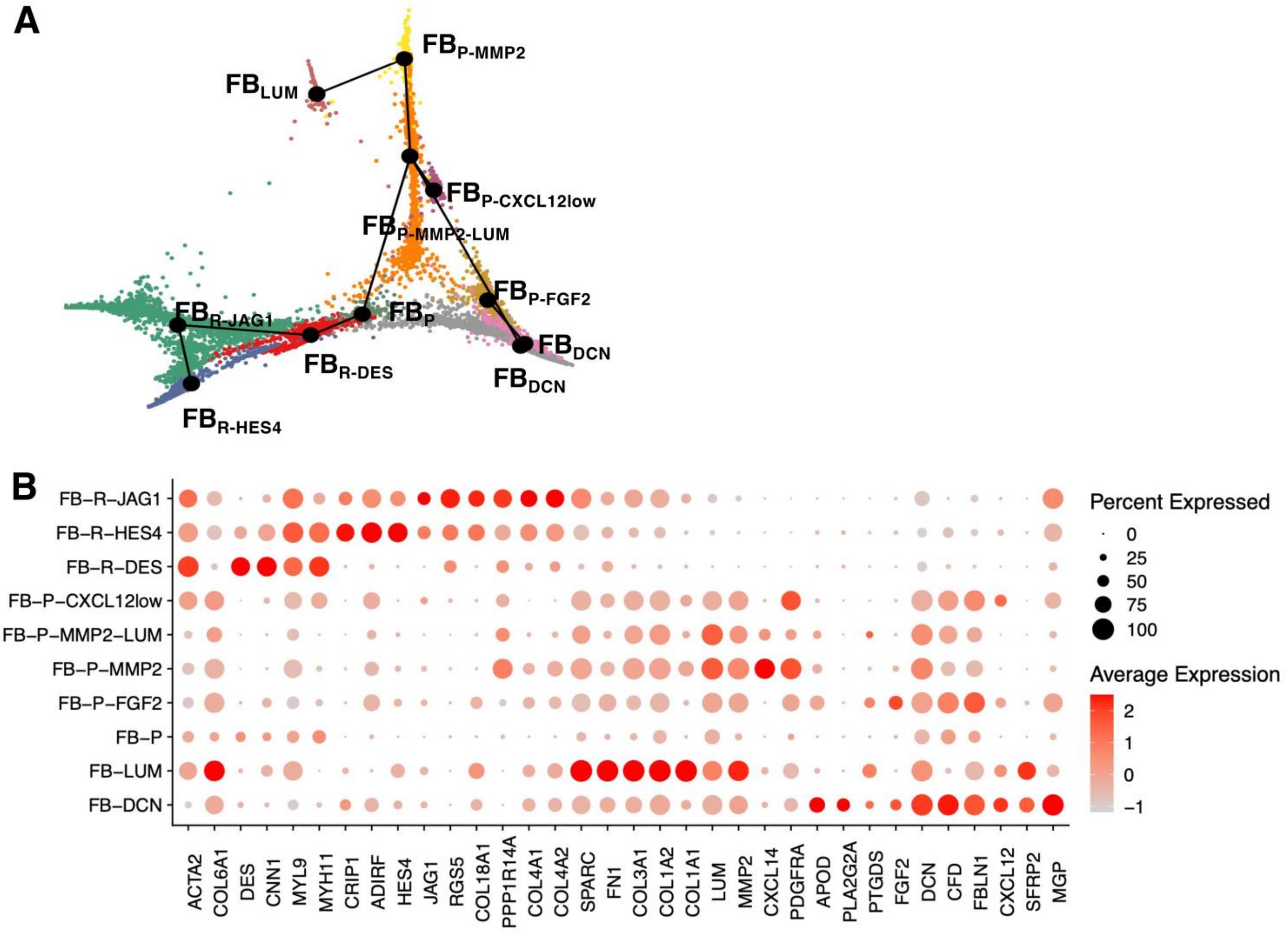


**Supplementary Figure 12. Fibroblast scRNA analysis.**

**(A)** Diffusion map projection overlaid with slingshot trajectory of single cell RNA sequencing with fibroblast cells from tissue. P-: PDGFRA+ve; R-: RGS5+ve. **(B)** Marker gene expression in different fibroblast clusters.

#### Supplementary Tables

**Supplementary Table 1. Human biospecimen and experiment details.**

**Supplementary Table 2. MIBC associated DMR loci.**

**Supplementary Table 3. scRNA marker genes.**

**Supplementary Table 4. scATAC marker peaks.**

**Supplementary Table 5. Cell Annotations for scRNA.**

**Supplementary Table 6. Cell Annotations for scATAC epithelial cells.**

**Supplementary Table 7. cisTarget-inferred TF-to-gene matches for NMF co-regulatory gene network in TPCS.**

**Supplementary Table 8. Cut&Tag peaks in RT4 and 5637 cells.**

**Supplementary Table 9. CNV of sequenced tumours.**

**Supplementary Table 10. SNV of sequenced tumours.**
